# Altered oral microbiota of drug-resistant organism carriers exhibit impaired gram-negative pathogen inhibition

**DOI:** 10.1101/2024.09.24.614756

**Authors:** Susan Zelasko, Mary-Hannah Swaney, Won Se Suh, Shelby Sandstrom, Caitlin Carlson, Julian Cagnazzo, Athena Golfinos, Jen Fossen, David Andes, Lindsay R. Kalan, Nasia Safdar, Cameron R. Currie

## Abstract

The oral microbiome has been understudied as a reservoir for clinical pathogens, including drug-resistant strains. Understanding how alterations in microbiome functioning render this site vulnerable to colonization is essential, as multidrug-resistant organisms (MDRO) carriage is a major risk factor for developing serious infections. To advance our knowledge of oral MDRO carriage and protection against pathogen colonization conferred by native microbiota, we examined microbiomes from individuals colonized by MDROs (n=33) and non-colonized age-matched controls (n=30). Shotgun metagenomic analyses of oral swabs from study participants revealed significant differences in microbial communities with depletion of *Streptococcus* spp. among those colonized by multidrug-resistant gram-negative bacilli (RGNB), compared to non-carriers. We utilized metagenomic sequencing to characterize the oral resistome and find antimicrobial resistance genes are present in higher abundance among RNGB carriers versus non-carriers. High-throughput co-culture screening revealed oral bacteria isolated from MDRO non-carriers demonstrate greater inhibition of gram-negative pathogens, compared to isolates from carriers. Moreover, biosynthetic gene clusters from streptococci are found in higher abundance from non-carrier microbiomes, compared to RGNB carrier microbiomes. Bioactivity-guided fractionation of extracts from *Streptococcus* isolate SID2657 demonstrated evidence of strong *E. coli* and *A. baumannii* inhibition in a murine model of infection. Together, this provides evidence that oral microbiota shape this dynamic microbial community and may serve as an untapped source for much-needed antimicrobial small-molecules.

## Introduction

Pathogens evolving to become multidrug-resistant are projected to give rise to over 10 million antibiotic resistant infections annually within the next 30 years [1, 2]. Without new and effective antibiotics, many life-saving medical procedures and immunosuppressive therapies will become too dangerous to perform. Diminishing antimicrobial drug development over the last forty years, owing to high rates of compound re-discovery, has been further complicated by the emergence of multidrug-resistance (MDR) owing to extreme selective pressure from antibiotic use in medicine and agriculture. Infection by MDR bacteria can be fatal in healthy individuals, and poses still greater risk for vulnerable populations such as the elderly, immunocompromised, and residents of regions with suboptimal healthcare infrastructure [3–6]. Moreover, MDR infections are often preceded by skin or mucosal colonization for variable lengths of time, resulting in asymptomatic inter-individual pathogen spread [7–9].

The oral microbiome is the second only to the gut microbiome in terms microbial abundance and diversity[10]. Oral microbes are the primary source of lung microbiota in adults, serving as gatekeepers between the external environment and lower aerodigestive tracts [11]. Antimicrobial resistances genes (AMRs) within pathogenic and commensal species, or the so-called “resistome,” have been previously characterized within oral microbiomes and shown to undergo frequent horizontal gene transfers [12–14]. However, oral carriage of drug-resistant organisms and the resulting implications for host health remain poorly understood. Prior investigations have primarily examined hospitalized populations [15–17], while carriage rates of other human pathogens among community-dwelling, adult populations remain less well known.

In defense against colonization, human-associated microbes are known to inhibit pathogens through various mechanisms, including production of antimicrobial specialized metabolites. For instance, nasal-associated *Staphylococcus lugdunensis* produces lugdunin, a cyclic nonribosomal peptide with bactericidal activity against vancomycin-resistant *Enterococcus* [18, 19]. Comparison of BGCs distribution across body sites reveals differential genomic capacity to synthesize ASMs, with the typical oral cavity containing relatively high BGC abundance [20]. As this site is in frequent contact with the environment and invading pathogens, oral microbiota appear to utilize ASMs in defense against pathogen spread to the lower gastrointestinal and respiratory tracts. Yet, functional characterization of intermicrobial interactions occurring via ASMs remains limited, especially among oral microbiota.

Here, we examine the oral microbiomes of adult multidrug-resistant organism carriers and non-carriers, previously characterized for presence of methicillin resistant S. aureus (MRSA), vancomycin-resistant *Enterococcus* (VRE), and resistant gram-negative bacilli (RGNB) through the Winning the War on Antibiotic Resistance (WARRIOR) study [21]. We utilize shotgun metagenomic sequencing to examine differences in composition, alpha diversity, antimicrobial resistance gene content, and biosynthetic gene cluster abundance from key oral taxa. High-throughput co-culture assays are used to compare inhibition of clinically-relevant pathogens by oral and fecal isolates from MDRO carriers versus non-carriers. Lastly, chemical extracts from a priority oral strain are evaluated for pathogen inhibition in a murine model of *E. coli* and *A. baumannii* infection. This study advances our understanding of oral drug-resistant pathogen carriage and resistomes, along with human-associated microbes contributing to pathogen inhibition through antibacterial small-molecule production.

## Results

### Composition of oral microbiomes from MDRO carriers and non-carriers

We obtained an average of 42,846,350 paired sequence reads from oral samples (n=63). After quality-filtering, including removal of reads mapping to human genomes, we retained an average of 23,600,993 (range: 333,062 – 70,933,558 sequences) high-quality sequences. After additional filtering, including via the decontam R package, oral metagenomes consisted of 4,803 bacterial species across 40 phyla and 56 fungal species across the phyla *Ascomycota* and *Basidiomycota,* in line with prior metagenomic studies of oral microbiomes [22]. Among the oral samples evaluated through the WARRIOR study (n=876), 9, 11, and 13 were found to be oral carriers of MRSA, VRE, and RGNB, respectively. The present study also evaluated 30 oral microbiomes from MDRO non-carriers. The most abundant genera from MDRO non-carriers, as well as MRSA and VRE carriers, included *Streptococcus, Prevotella, Veillonella, Rothia*, and *Neisseria* (Figure 1A). Among RGNB carriers, the top genera included *Streptococcus, Prevotella, Veillonella, Acinetobacter*, and *Pseudomonas* (Figure 1A).

**Figure 1.**
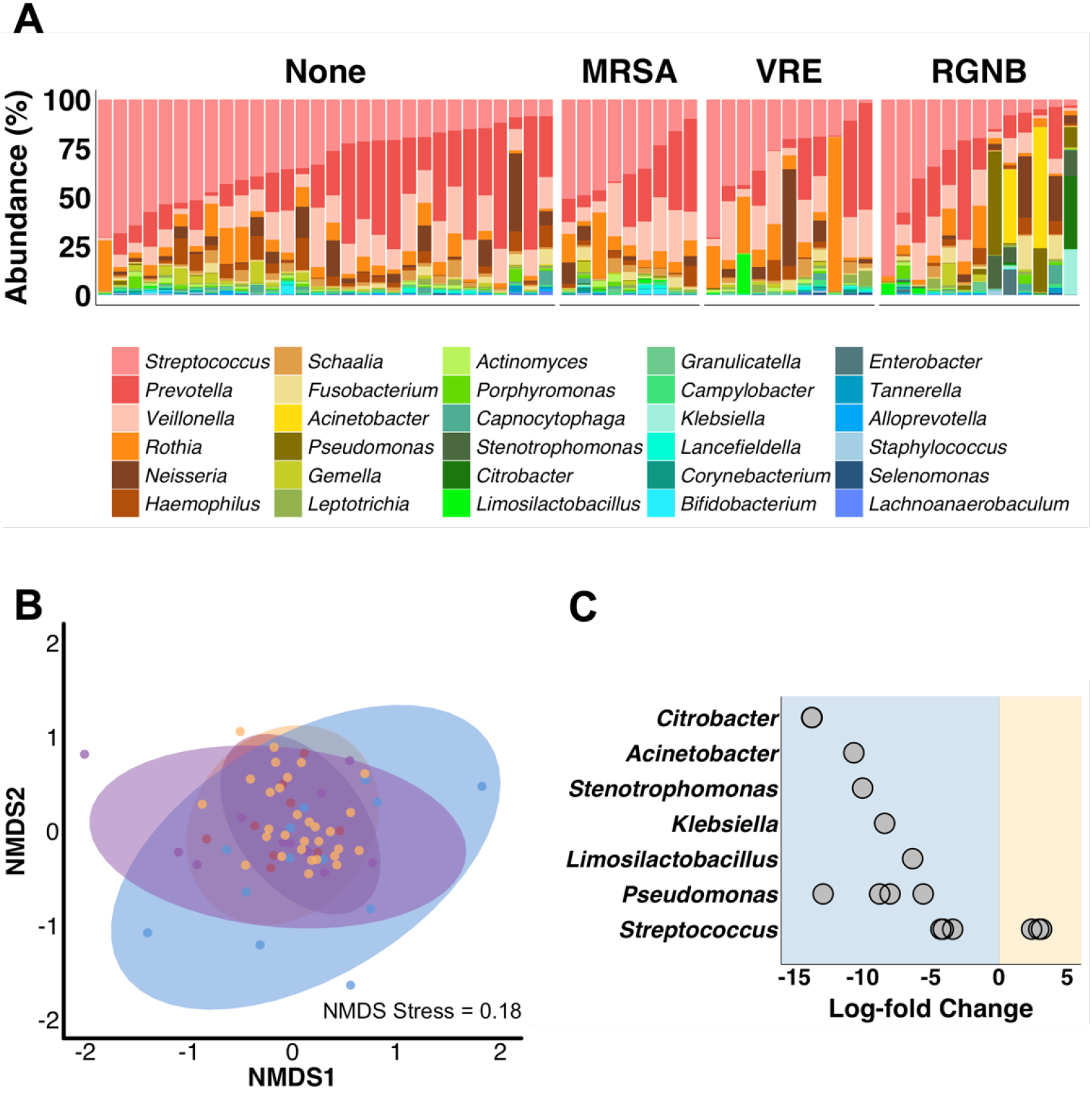
Oral Microbiome Composition Varies Among MDRO Non-carriers and Carriers. (A) Relative abundance of top 30 genera grouped by MDRO non-carriers (“None”), MRSA, VRE, and RGNB carriers. (B) Non-metric multidimentional scaling (NMDS) ordination plot of Bray-Curtis dissimilarity indices of oral microbiomes from MDRO non-carriers (“None”), MRSA, VRE, and RGNB carriers, colored per legend. Points represent individual samples. (C) Plot shows differentially abundant taxa between MRDO non-carriers (yellow shaded) and RGNB carriers (blue shaded). Taxa with FDR corrected p-values < 0.05 displayed. Points represent individual species, grouped by genera on y-axis. For all analyses, shotgun metagenomic sequencing reads were examined from microbiomes normalized to an equal sequencing depth.

We next examined the impact of MDRO carriage on oral microbial community structure. Non-metric multidimentional scaling (NMDS) of Bray-Curtis dissimilarity values identified the microbial composition of oral microbiomes from MDRO non-carriers to be significantly different from those with RGNB (ANOSIM R=0.299, p.adj<0.01) (Figure 1B, Table S1). Comparisons of MRSA or VRE carriers relative to non-carriers did not reveal significant differences in beta-diversity (Table S1). No significant differences in richness, Shannon, Inverse Simpson, or Fisher alpha diversity metrics were identified across MDRO carriers or non-carriers, though there were trends toward higher diversity among microbiomes of MDRO non-carriers (Figure S1).

Differential abundance analysis comparing MDRO non-carriers to RGNB further revealed oral species enriched across each group. In total, 14 species were present in higher abundance among RGNB carriers, including *Stenotrophomonas maltophilia*, *Citrobacter spp.*, *P. aeruginosa*, *K. pneumoniae*, *Acinetobacter johnsonii*, as well as *Limosilactobacillus fermentum* and three *Streptococcus* spp. (FDR<0.05, Figure 1C, Table S2). In contrast, MDRO non-carrier oral metagenomes had higher abundance of three species of *Streptococcus* spp. (FDR<0.05, Figure 1C, Table S2).

### Detection of antimicrobial resistance genes from oral metagenomes

We next sought to define antimicrobial resistance (AMR) gene content and abundance across the oral microbiomes of MDRO carriers, relative to non-carriers. We subsampled metagenomes to a set read depth and aligned the sequences against the curated Comprehensive Antibiotic Resistance Database (CARD). We observed a set of 18 AMRs that were present in >70% of samples (Figure 2A, Figure S2, Table S3), 17 of which were present in >70% of MDRO non-carrier samples. Of these genes, 6 belonged to “tet family” conferring tetracycline resistance and 2 genes, tet(M) and tetB(46), were present in 100% of examined samples. In general, samples clustered by carriage or non-carriage of MDROs, which revealed a cluster of RGNB containing AMR genes from *P. aeruginosa, Stenotrophomonas maltophilia, E. coli, K. pneumoniae*, and *A. baumannii*. Accordingly, we identified higher median numbers of FPKM reads mapping to protein-coding AMR genes from RGNB carriers compared to non-carriers that approached statistical significance (91 versus 69, respectively, p = 0.056, Figure 2B). Metagenomes originating from carriers of VRE had lower AMR gene abundance, relative to non-carriers (57 versus 69, respectively, p <0.05, Figure 2B). No significant differences were identified between non-carriers and MRSA carriers. We further identified 120 and only 6 AMR genes that were differentially abundant in RGNB carriers versus non-carriers, respectively, while 90 were shared (Figure 2C, Table S4). Among RGNB carrier metagenomes, the AMR genes adeJ, oqxB, and acrB were the most significantly different from non-carrier metagenomes. Accordingly, the prevalence of these genes in RGNB carriers was higher compared to non-carriers (adeJ 23% vs. 3.4%; oqxB 38.5% vs. 0%; acrB 46.2% vs. 3.4%).

**Figure 2.**
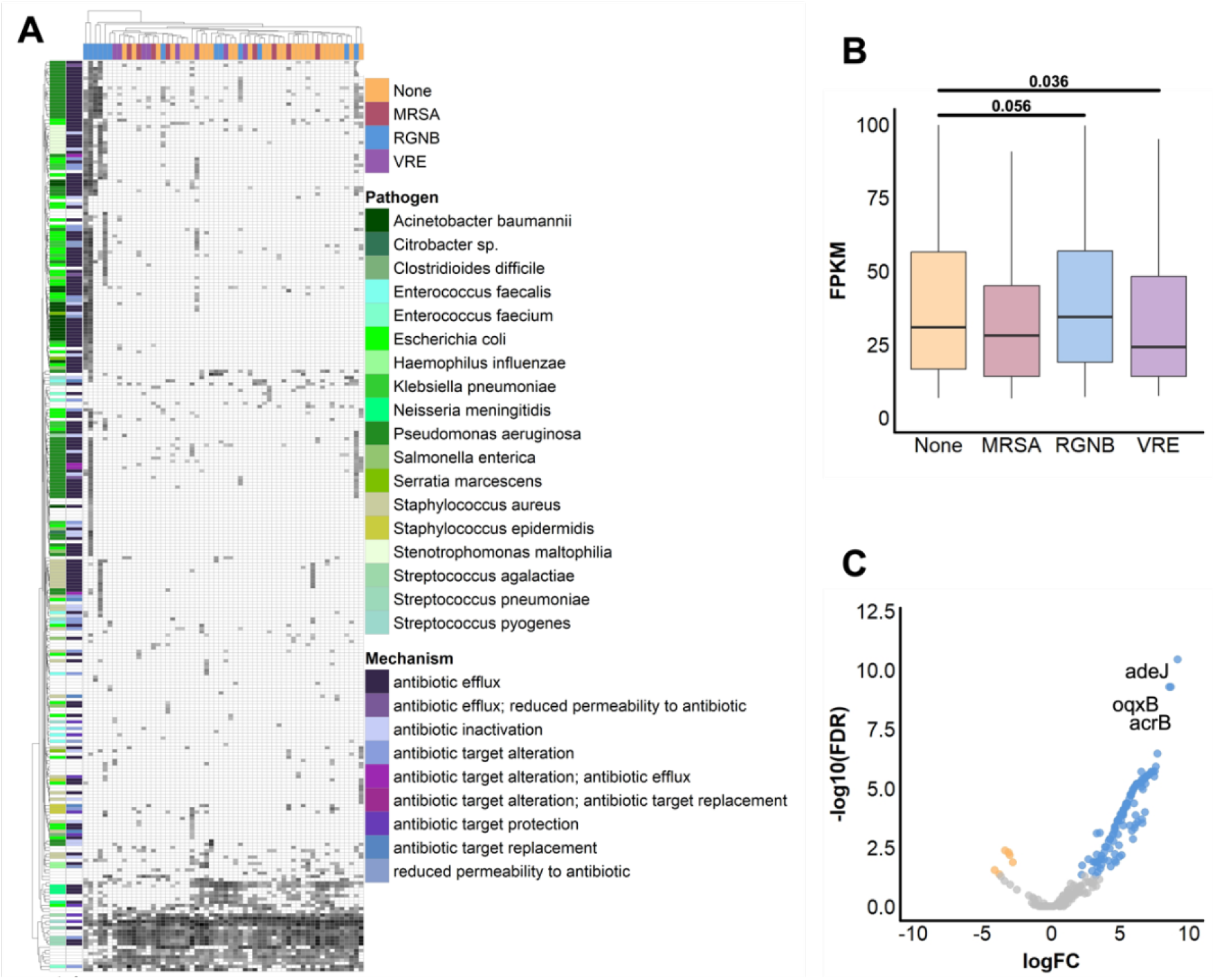
MDRO Non-carriers and Carriers Posses Differing Oral Resistomes. Metagenomes (n=58) were subsampled to 1,000,000 paired reads and aligned to the CARD database. (A) Heatmap displays log10 + 1 values of FPKM aligning to AMRs with MAPQ ≥ 10 (rows) across oral metagenomes from MDRO non-carriers and carriers (columns), colored per legend. Both rows and columns are hierarchically clustered. Dark grey values indicate higher FPKM values. Pathogen of origin and resistance mechanisms for each AMR are indicated on the left side of the plot, colored per legend. (B) Boxplots of FPKM aligning to antimicrobial resistance genes with MAPQ ≥ 10 between MDRO non-carriers and carriers. Boxes show the 25th and 75th percentiles with the median represented by a horizontal line and whiskers showing 1.5 x the interquartile range. Significance values from Mann–Whitney U-test are listed. (C) Volcano plot comparing AMRs identified from oral metagenomes that are differentially abundant between MDRO non-carriers (yellow points) and RGNB carriers (blue points), or not significantly different (grey points). Y-axis represents inverse of log of FDR significance values and x-axis represents the log fold-change between groups. FDR cutoff of <0.05 was used. Select genes are labeled.

### Oral microbiota of MDRO non-carriers inhibit gram-negative pathogens

Having identified significant enrichment of streptococci in oral microbiome of MDRO non-carriers, relative to RGNB carriers, we sought to examine pathogen inhibition by microbiota across these groups. We hypothesized that isolates of *Streptococcus* originating from MDRO non-carriers contribute to inhibition of gram-negative pathogens. To this end, we performed high-throughput co-culture assays of bacteria isolated from WARRIOR microbiomes samples [20]. We screened 152 (n=31 hosts) and 140 (n=35 hosts) oral and fecal isolates, respectively, against seven gram-positive, nine gram-negative, and five fungal pathogens. In total, 6,841 pairwise interactions were evaluated. Comparison of fractional inhibition identified that gram-negative pathogen inhibition was significantly higher among oral isolates originating from MDRO non-carriers, relative to carriers (0.063±0.0081 vs. 0.025±0.0075, p.adj <0.05 Figure 3, Table S5). No significant differences in fractional inhibition of fungal or gram-positive pathogens by oral isolates from carriers versus non-carriers were identified, nor were there significant differences in rates across pathogens types from fecal isolates.

**Figure 3.**
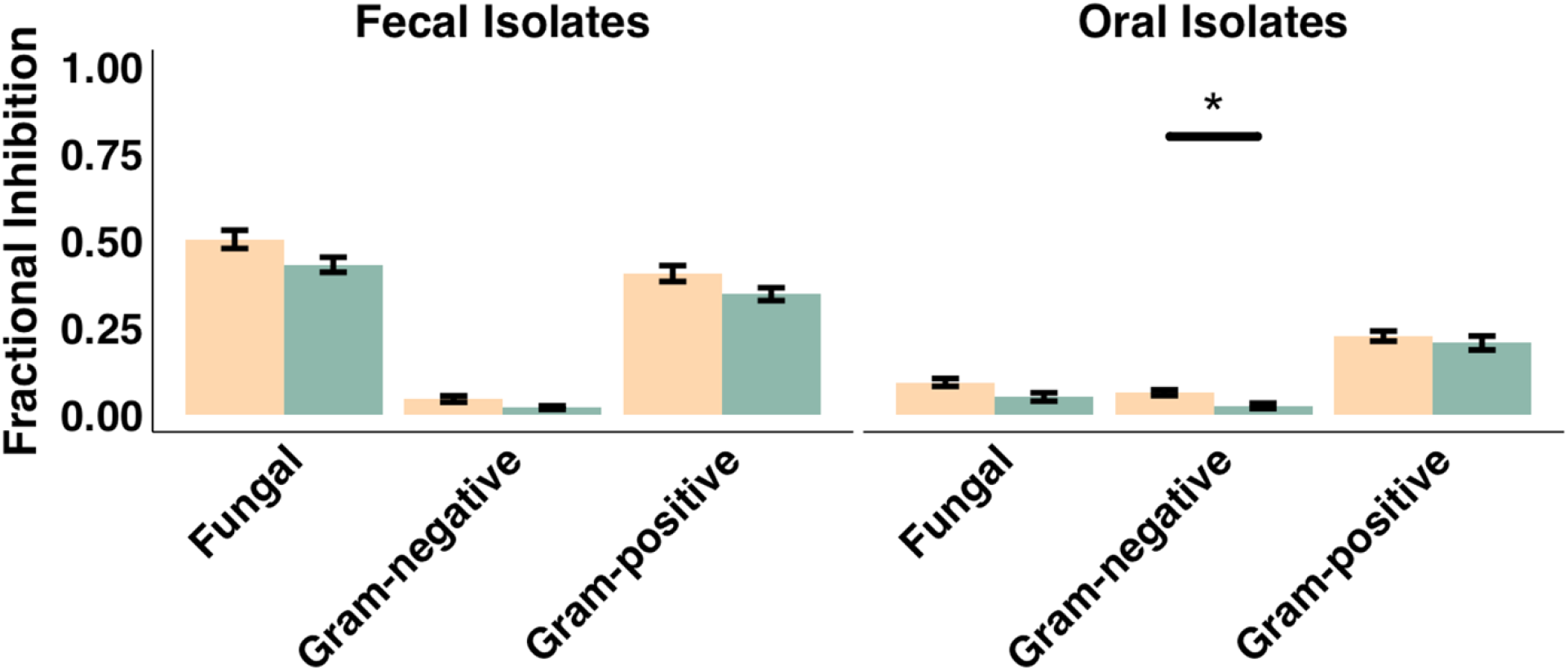
Contact-independent Pathogen Inhibition by Isolates from MRDO Carriers compared to Non-carriers. Plot displays fractional inhibition scores from co-culture bioactivity assays for fecal and oral isolates across fungal, gram-negative, and gram-positive pathogens. Significance value * = p < 0.05 Mann– Whitney U-test.

### MDRO Non-carriers have Higher BGC Abundance relative to MDRO Carriers

We next sought to evaluate BGC abundance across microbiomes of MDRO non-carriers and carriers to examine how specialized metabolite production may contribute to differing rates of pathogen inhibition. We first examined the abundances of subsampled metagenomic reads aligning to non-ribosomal polyketide synthetase (NRPS), polyketide synthase (PKS), ribosomally synthesized and post-translationally modified peptide (RiPP), terpene, or hybrid BGCs originating from oral- and nasal-associated bacteria (Figure 4A). A greater mean number of reads from oral microbiomes of MDRO non-carriers mapped to NRPS, PKS, and RiPP BGC open reading frames, relative to RGNB carriers (Figure 4A). MDRO non-carriers also had higher mean RiPP BGCs relative to VRE carriers and higher terpene BGCs relative to MRSA carriers. Next, we examined the abundances of these clusters across the top five most abundant oral genera (Figure 4B). We found MDRO non-carriers had higher mean abundance of NRPS, RiPP, and hybrid clusters originating from streptococci, relative to RGNB and VRE carriers. Additional differences in mean abundances of terpenes from *Neisseria*, PKS from *Prevotella*, RiPPs from *Veillonella*, and hybrid clusters from *Prevotella* and *Rothia* were noted between MDRO non-carriers and carriers.

**Figure 4.**
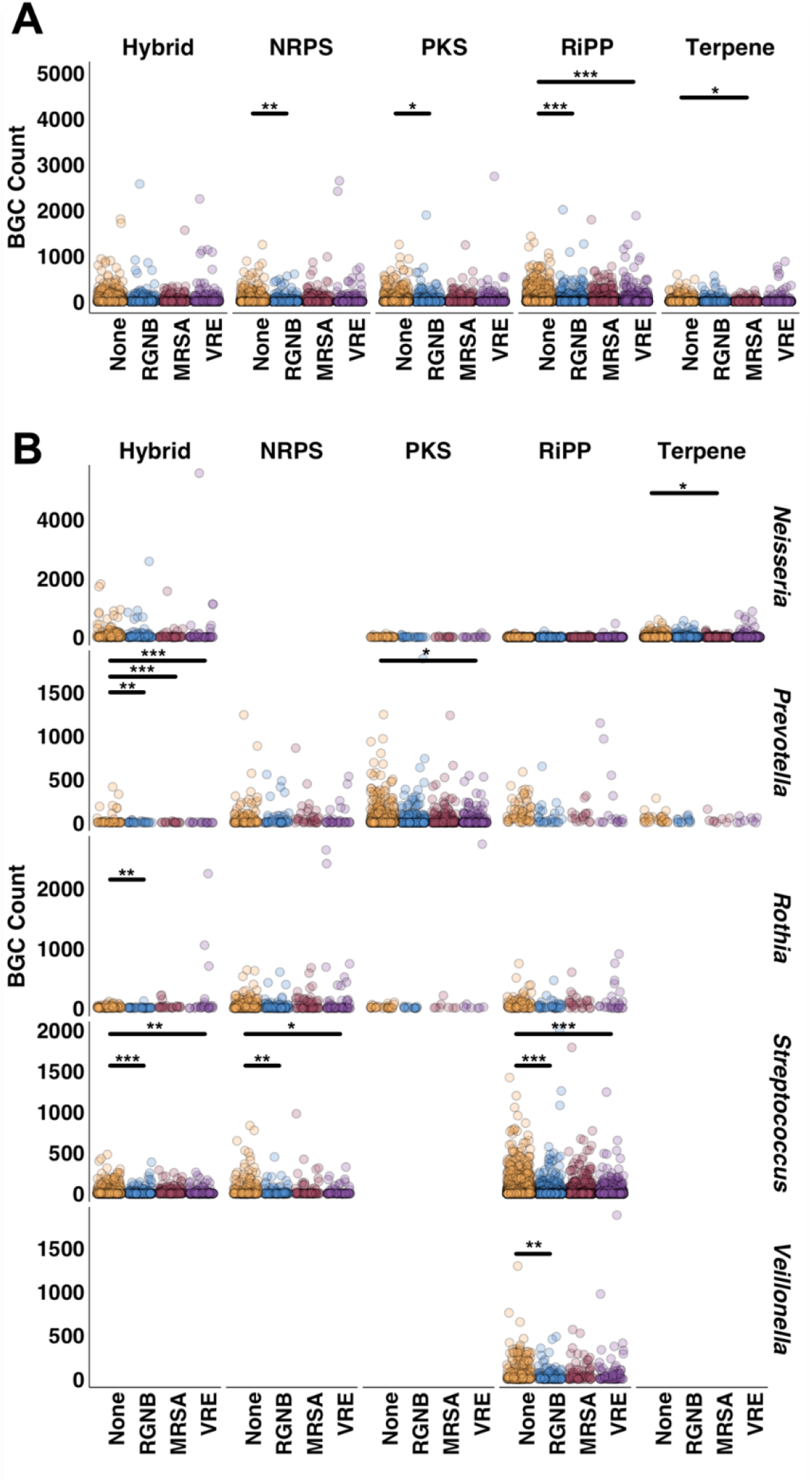
MDRO Non-carriers have Higher BGC Abundance relative to MDRO Carriers. (A) The plot indicates reads from nasal (blue) and oral (red) infant and adult microbiomes that mapped to BGCs (arylpolyene, NRPS, PKS, RiPP, siderophore, terpene, and hybrid clusters) identified from upper respiratory bacteria (eHOMD habitat listed as “Nasal”, “Nasal,Oral”, or “Oral”). (B) Reads from nasal (blue) and oral (red) infant and adult microbiomes mapping to BGCs from upper respiratory bacteria, grouped by cluster type. All metagenomes were subsampled to 160K reads. 51 infant nasal, 22 adult nasal and 1 adult oral metagenomes were not included in analysis due to low sequencing depth. Normalized reads were pseudoaligned onto an in-house BGC library. NRPS = non-ribosomal peptide synthases, PKS = polyketide synthases, RiPP = ribosomally synthesized and post-translationally modified peptides. Significance value * = p < 0.05, ** = p <0.01, *** = p <0.001, **** = p <0.0001 Yuen Welch’s T-test with 0.001 trim.

### Fractionated extract of *Streptococcus* SID2657 inhibits *E. coli* and *A. baumanii* in a murine model of infection

We next investigated the contact-independent inhibition mediated by isolate SID2657 against gram-negative pathogens. SID2657 was grown in liquid media with Diaion HP20 resin, yielding 67.5 mg/L of ethyl acetate crude extract containing metabolites produced by this strain. Following fractionation of crude extract by a flash chromatography, all fractions were screened for pathogen inhibition via disc diffusion. One fraction exhibited inhibition of *E. coli* and *A. baumannii*, while another fraction inhibited *S. aureus* (Figure 5A). No fractions inhibited *C. albicans*. Next, extract fractions were evaluated for gram-negative pathogen inhibition in a murine model of infection (Figure 5B). Mice treated with the fraction of interest had 1.25 and 0.97 log-reductions in infectious burden of *E. coli* and *A. baumannii* with a single dose, respectively (Figure 5C).

**Figure 5.**
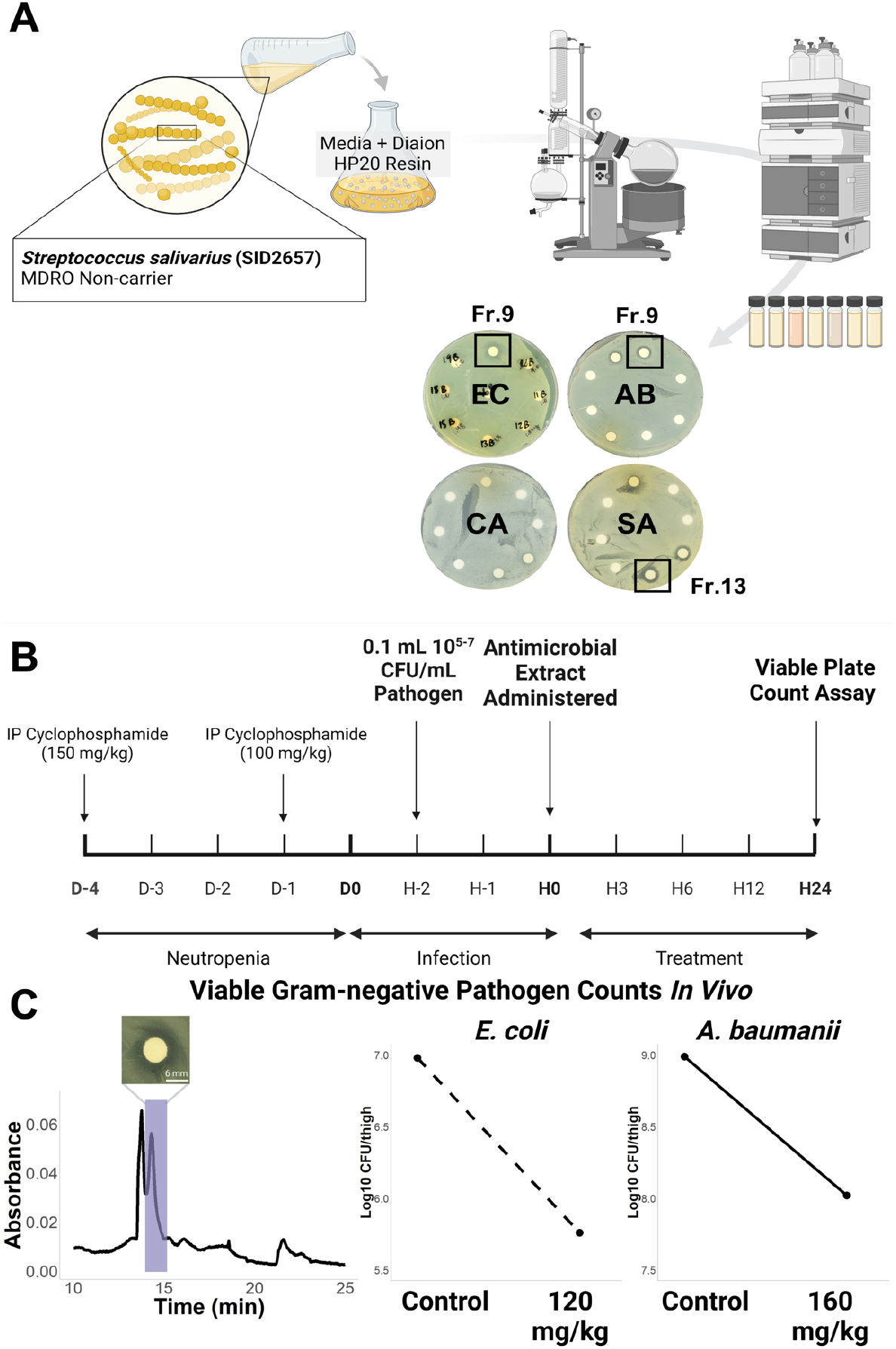
Extract Fraction from *Streptococcus* sp. Inhibits Gram-negative Pathogens in an *In Vivo* Infection Model. (A) *Streptococcus* SID2657 was inoculated 1:1000 in Basic Medium with Dianon HP20 Resin and incubated at 37C for 6 days. Washed resin was extracted with an equal volume of ethyl acetate (EA) and dried, then fractionated by flash chromatography using stepwise MeOH/H2O gradient. Overnight cultures of *E. coli, A. baumannii, C. albicans*, and *S. aureus* were adjusted to 0.5 McFarland standard and plated onto Mueller-Hinton agar. Blank discs were spotted with extract fractions dissolved in MeOH (2 mg/disc) and placed onto plates. Representative disc diffusion assays are shown. (B) Timeline for murine mouse model experiments is displayed. D: days, H: hours. (C) Chromatograph of crude extract fractionation is shown (left) with highlighted peak corresponding to fraction with gram-negative activity, as demonstrated by representative disc diffusion assay image. Log of CFU/thigh from viable plate count assays for *E. coli*- and *A. baumannii*-infected mice are shown following control vehicle and extract fraction administration.

## Discussion

To our knowledge, this study represents the first metagenomic comparison of oral microbiomes across healthy, community-dwelling adults who are MDRO carriers versus non-carriers. While there has been growing recognition that the oral cavity serves as a reservoir of pathogens and AMRs [12, 23], few works have examined how commensal microbiota may prevent pathogen colonization at this site. To this end, our study uses shotgun metagenomic sequencing to identify key bacteria present in higher abundance among MDRO non-carrier microbiomes and demonstrates their capacity to inhibit gram-negative pathogens.

First, our data identify that oral microbial community composition is significantly disrupted among RGNB carriers, which contain greater abundance of gram-negative taxa and lower abundance of specific streptococci relative to non-carriers. Limited prior studies have identified gram-negative bacilli to be present as non-core members of the oral cavity, primarily among those with poor oral health, systemic medical conditions, and who are hospitalized or reside in long-term care facilities [16, 17, 24–26]. We find RGNB carrier microbiomes contained known pathogens (*P. aeruginosa, K. pneumoniae, Stenotrophomonas maltophilia*), as well as the putative pathogen *Acinetobacter johnsonii* that has been noted to encode several AMR genes [27, 28]. The implications of gram-negative carriage include predisposition toward respiratory infection by these pathogens [29, 30]. In contrast to RGNB carriers, no significant differences in community composition we identified among MRSA and VRE carriers, possibly owing to low abundance carriage of these pathogens.

We next examined oral resistomes across groups, finding both AMR genes common to the majority of samples and a subset that are unique to RGNB carriers. The oral microbiome has been increasingly recognized as a reservoir of AMR genes. Here, we show that MDRO non-carriers possess a median of 69 AMR genes and that 18 AMRs are present in over 70% of all examined samples. These prevalent AMRs confer resistance to a range of antibiotics, including tetracyclines, macrolides, and fluoroquinolones, and corroborate results of prior studies [12, 31]. The finding of common AMR genes across populations, including remote-dwelling and ancient humans, suggests components of the resistome are natural features of the oral microbial communities [32, 33]. We further show a trend toward higher AMR abundance among RGNB carriers, particularly samples dominated by *Acinetobacter*, *Citrobacter*, and *Pseudomonas*, with enrichment of a subset of 106 resistance genes relative to non-carrier microbiomes. This included many genes conferring multidrug resistance through efflux pump mechanisms. Carriage of pathogens with AMR genes constitutes a reservoir for subsequent colonization of other sites and for transmission to others, promoting further dissemination of challenging to treat infections.

Despite its clear association with a range of diseased or healthy states, the role of oral microbiota in colonization resistance has remain incompletely understood. The observed dysbiosis of RGNB carrier oral microbiomes, relative to non-carriers, suggested an altered capacity of resident taxa to engage in defense against colonization by pathogens. We therefore compared rates of pathogen inhibition by commensal isolates via high-throughput co-culture assays. Frequency of gram-negative pathogen inhibition was overall low, in line with many intrinsic and acquired defenses such pathogens have to inhibition [34, 35]. However, oral isolates of MDRO non-carriers exhibited higher fractional inhibition of gram-negative pathogens relative to carriers. Fecal isolates followed a similar pattern that did not achieve significance, but may point toward a wider alteration of microbiota among MDRO carriers across body sites.

Next, we corroborated differences in contact-independent inhibition of gram-negative pathogens by examining BGC abundances across oral microbiomes. BGCs encode for the production of diverse metabolites that enable microbial niche-adaption and interactions with other microbiota and the host [20, 36]. Of note, many specialized metabolites produced by NRPS, PKS, RiPP and some terpene or hybrid clusters possess antimicrobial activity [37]. We demonstrate that across oral microbiota, MDRO non-carriers have a higher mean number of BGCs belonging to NRPS, PKS, and RiPP classes. Examination of patterns of BGCs across major oral taxa further show streptococci from non-carriers have higher NRPS, RiPP, and hybrid abundance. Examination of BGC abundance across other major taxa revealed that *Prevotella* and *Rothia* hybrid clusters, as well as RiPPs from *Veillonella* were also more abundant among MDRO non-carriers compared to RGNB carriers. Together, this suggests a range of bacteria inhabiting MRDO non-carriers possess greater capacity for production of specialized metabolites with potential to that modulate microbe-microbe and microbe-host interactions. Oral streptococci have previously been demonstrated to produce antimicrobial small-molecules [38, 39], while the functions of hybrid- and RiPP-encoded metabolites from non-streptococcal taxa remain unstudied.

To begin to examine the functional role of specialized metabolites produced by oral microbes, we examined bioactivity of a priority *Streptococcus* sp. Whole-genome sequencing of this strain identified novel RiPP and type III PKS clusters that did not encode predicted or known metabolites. We show that extracted metabolites from this isolate exhibited *in vitro* gram-negative pathogen inhibition was shown to inhibit *E. coli* and *A. baumannii* in a murine model of infection. Prior investigations have identified oral streptococci that demonstrate contact-independent inhibition of *P. aeruginosa* via hydrogen peroxide and nitrite production [40], as well as respiratory pathogens *M. catarrhalis* and *S. pneumoniae* [41]. Neither of these mechanisms mediated observed interactions herein, as these labile compounds would not be extracted by ethyl acetate. Some oral streptococci produce RiPP-encoded bacteriocins, however these molecules are not reported to possess anti-gram-negative activity [39], nor were any known bacteriocins predicted in the genome of the isolate examined. More broadly, antagonistic relationships between *Streptococcus* spp. and other gram-negative pathogens have, until this work, not been reported. Oral streptococci are key members of polymicrobial biofilms that serve as the interface between host mucosa, immune cells, and the bloodstream [42, 43]. Non-oral species, including *E. coli*, have been shown to incorporate into dental biofilms [44], providing potential means of accessing systematic circulation or the gut. Translocation of oral bacteria to arterial plaque and blood has been well-documented and linked with a range of systemic inflammatory diseases [45, 46]. Enteral accumulation of oral *Enterobacteriaceae* has likewise been noted in the setting of periodontal inflammation [25]. Thus, it is conceivable that commensal streptococci may utilize diffusible small-molecule mediators to prevent exogenous gram-negative microbes from integrating into the shared biofilm niche, thereby maintaining host health.

In summary, we examine for the first time the oral microbiomes of MDRO carriers and non-carriers using shotgun metagenomics and functional bioassays. We show microbial communities of RGNB carriers significantly differs in terms of composition and AMR gene content from that of non-carrier microbiomes. Oral bacteria isolated from MDRO non-carriers demonstrated greater inhibition of gram-negative pathogens, suggesting that MDR carriage-associated dysbiosis is accompanied by loss of protective species. Accordingly, we find enrichment of *Streptococcus* spp. among non-carriers and show metabolites from a priority isolate inhibit *E. coli* and *A. baumannii* in a murine model of infection. This adds to growing evidence that microbial competition in the oral microbiome serves as a first line of defense against pathogen colonization, with important implications for host infection risk.

## Materials and Methods

### Study Design and Sample Collection

Biospecimen samples (saliva, oral swabs, stool) were collected through the WARRIOR study [21], as an ancillary study to the Survey of the Health of Wisconsin (SHOW), an ongoing cross-sectional survey of residents across Wisconsin. SHOW participants age >18 (see published exclusion criteria[21]) were invited to participate in the WARRIOR study (n=876 participants). From 2016-17, saliva (1-2 mL) was collected into a sterile tube, buccal mucosa/tonsils were swabbed via sterile BBL CultureSwab with liquid Stuart transport medium (Becton, Dickinson and Company, Franklin Lakes, New Jersey, USA) during study clinic visits, and stool was self-collected into collection kits. All samples were received within 24 hours by the UW-Madison Infectious Disease Research Lab for multidrug-resistant organism (MDRO) colonization testing, followed by storage at -80°C, as previously described [21]. The present study performed metagenomic sequencing analyses using all oral samples with any detected MDRO (n=33 participants), along with age-matched control samples without MDROs (n=30 participants). For co-culture experiments, salivary and fecal samples from MDRO carriers (n=12 and n=18 participants, respectively) and non-carriers (n=19 and n=17 participants, respectively) were used. The WARRIOR study was approved by the Human Subjects Committee at the University of Wisconsin-Madison.

### DNA extraction and Metagenomic Sequencing

DNA extractions of oral swab samples were carried in a DNAse, UV-treated, and bleach-treated laminar flow hood. Negative DNA extraction controls (n=8) were processed and sequenced in tandem with study samples. Two mock microbial communities contained either 20 Strain Staggered Mix Genomic Material (ATCC MSA-1003) or 6 Strain Skin Microbiome Genomic Mix (ATCC MSA-1005) were also included as positive controls. Prior to extraction, samples were thawed on ice and 100 μL of storage media was combined with ReadyLyse (Epicenter), Mutanolysin (Sigma), and Lysostaphin (Sigma), followed by incubation at 37°C with shaking for 1 h. Samples were next added to a glass bead tube (Qiagen) and vortexed for 10 min followed by incubation at 65°C with shaking for 30 min. After transferring sample liquid to a sterile tube, MPC protein precipitation reagent (Lucigen) was added. The sample was vortexed and centrifuged for 10 min at 21,300 × *g*. The supernatant was combined with isopropyl alcohol and column purified (Invitrogen PureLink Genomic DNA Extraction Kit) then eluted in 50 μL elution buffer. DNA was quantified using the PicoGreen dsDNA quantification assay kit (Invitrogen). Illumina sequence libraries were prepared using the Nextera XT DNA Library Preparation Kit (Illumina), followed by pooling and sequencing at the University of Minnesota Genomics Center. 2×150 bp paired-end sequencing of pooled samples was performed on the NovaSeq 6000.

### Metagenomic Sequence Processing

Raw read data was first preprocessed using fastp v0.21.0 [47] to filter low quality, reads <50 bp, and remove adapters sequences. Filtered reads were processed using Kneaddata v0.8.0 to remove tandem repeats using Tandem Repeat Finder and to filter and remove human DNA sequences by mapping reads to the human genome. Taxonomic identification was assigned using Kraken2 v2.0.8-beta [48] and abundances estimated using Bracken v2.5 [49], with a custom database including complete bacterial, viral, archaeal, fungal, protozoan, and human genomes along with UnivEc ore sequences from RefSeq. The database separated and filtered plasmid sequences, as plasmid sequences are frequently assigned incorrect taxonomies using Kraken2 [50]. Potential contaminant species were identified from negative extraction controls and excluded from further analysis using the prevalence method in decontam v1.10.0 (threshold = 0.25) [51]. Reads not annotated at the phylum level, as well as those belonging to phyla present at <1% prevalence were additionally filtered.

### Metagenomic Sequence Analyses and Statistics

Nonparametric multidimensional scaling (NMDS) was performed on Bray-Curtis dissimilarity values of normalized read counts using the ordinate function of the phyloseq R package (version 1.42.0) [52]. ANOSIM were performed on beta diversity distances with 999 permutations using the vegan R package (version 2.6-4) [53]. Alpha diversity metrics were determined by comparing normalized read counts using the estimate_richness function of the phyloseq R package (version 1.42.0) [52] and compared using the Mann–Whitney U-test. Mean read count pseudoaligned to BGCs was compared using Yuen’s T test with 0.001 trim.

### Detection and Analysis of Antimicrobial Resistance Genes

Processed metagenomic sequencing reads were subsampled to a set count (1 million reads) via seqtk 1.4. Four samples were removed from analysis due to sequencing depth <1M reads. Antibiotic resistance genes were identified by alignment of metagenomes against the Comprehensive Antibiotic Resistance Database (CARD) via KMA (v1.4.9) using RGI (v5.1.0) (rgi bwt -read_one -read_two -output_file -clean -local) [54]. Each gene was annotated with Resistance Mechanism and Pathogen of Origin using CARD metadata. The read count abundance was normalized to Fragments Per Kilobase Million FPKM by the formula: FPKM = (counts x 1E9) / (gene_length x sum(counts)), where “counts” represents the number of reads mapped, “gene_length” is the length of the corresponding gene. Median FPKM aligned to antimicrobial resistance genes were compared across groups using the using the Mann–Whitney U-test. Differential abundances of resistance genes were calculated using the edgeR package in R (v3.32.1) [55, 56].

### Alignment of Metagenomic Reads to Biosynthetic Gene Clusters

Processed metagenomic sequencing reads were subsampled to a set count (1 million reads) via seqtk 1.4. Kallisto quant 0.46.0 [57] was used to pseudoalign reads to an index of URT-associated BGC open reading frames (ORFs), as previously described [36]. Briefly, 1,527 bacterial genome sequences were downloaded from the eHOMD V9.03 [58], from which 3,895 BGCs were identified via antiSMASH [59]. Hmmscan from HMMER 3.1b2 [60] was then used to identify ORFs commonly associated with BGCs. Kallisto was used to build a 31 length k-mer index containing all ORFs, except nonbiosynthetic ORFs. All estimated read counts for all ORFs were aggregated into a single estimated read count per BGC.

### Strain Isolation and Identification

Sample storage medium (100 μL) was inoculated onto BHI agar and incubated for 5 days at 37°C. Colonies of distinct morphotype (≥2 colonies per plate) were selected from each sample and passaged until non-contaminated monocultures were obtained. Each isolate was inoculated in 3 mL of BHI and grown overnight at 37°C with shaking. All isolates were stored at −80°C in 20% (vol/vol) glycerol.

### Co-culture Inhibition Bioassays and Scoring

Pathogen inhibition of fecal and oral strains was assessed using co-culture plate inhibition assays, as previously described [61, 62]. Briefly, each isolate was inoculated in 3 mL of BHI and grown overnight at 37°C with shaking. Liquid cultures were inoculated onto one half of each well in a 12-well plate containing 3 mL/well of BHI. Plates were incubated at 37°C for 7 days. Bacterial and yeast pathogens were grown in 3 mL of BHI overnight at 28°C with shaking. Pathogen cultures were diluted 1:10 and 1 μL of diluted culture was spotted on the center of each well. Spore stocks of filamentous fungi (stored at −80°C) were diluted 1:10 in BHI prior to inoculation. Plates were again incubated at 28°C for 7 days. Pathogen inhibition was scored to 3 as follows: 0 – no inhibition, 1 – partial inhibition, 2 – presence of a zone of inhibition, 3 – total inhibition. Scores for wells with lack of isolate growth, contamination, or overgrowth were not determined. Fractional inhibition was calculated as previously described [61].

### Whole Genome Sequencing, Assembly, and BGC Detection

Isolate SID2657 was grown in BHI media with shaking for 18 hrs at 37°C. Cells were harvested by centrifuging the cultures at 21,330 × g for 5 min. DNA was extracted from cell pellets (Biosearch Technologies) and quantified using the Qubit dsDNA BR quantification assay kit (Invitrogen). Genomic libraries for Illumina MiSeq 2×150-bp paired-end sequencing were prepared and sequenced by SeqCenter (Pittsburgh, PA). Paired ends were assembled with SPAdes v3.11.0. and BGCs were identified within each genome with antiSMASH v7.0.

### Generation of Extract Fractions

Isolate SID2657 was grown in Basic Media[18] overnight at 37°C and used to inoculate large volume culture (1:1000) containing Diaion HP20 resin (10g/L). Cultures were grown at 37°C for 6 days with shaking. Filtered resin beads were separated from liquid media and washed with Milli-Q water to remove media components and primary metabolites, then incubated with ethyl acetate (v/v) overnight then dried by rotatory evaporation. Extract was fractionated by flash chromatography using a stepwise 25/50/75/100% MeOH/H_2_O gradient, with all peaks collected and dried by rotary evaporation.

### Disc Diffusion Assays

Overnight cultures of *E. coli, A. baumannii, S. aureus*, and *C. albicans* grown at 28°C were adjusted to 0.5 McFarland standard and plated onto Mueller-Hinton agar. Blank discs were spotted with fractions dissolved in MeOH (2 mg/disc), dried in ambient air, and placed onto plates, with methanol serving as a negative control and gentamicin, mupirocin, or amphotericin serving as positive controls for *E. coli* and *A. baumannii*, *S. aureus*, and *C. albicans*, respectively. Plates were incubated overnight at 37°C, followed by examination of zones of inhibition by extract fractions.

### Mouse bacterial thigh infection model

Animals for the present studies were maintained in accordance with the criteria of the Association for Assessment and Accreditation of Laboratory Animal Care. All animal studies were approved by the Animal Research Committee of the William S. Middleton Memorial VA Hospital. Experiments used six-week-old, specific-pathogen-free, female ICR/Swiss mice weighing 23–27 g (Harlan Sprague-Dawley, Indianapolis, IN). Mice were rendered neutropenic (neutrophil count <100 mm^−3^) via subcutaneous cyclophosphamide injection (Mead Johnson Pharmaceuticals, Evansville, IN) 4 days (150 mg/kg) and 1 day (100 mg/kg) prior to thigh infection. Broth cultures of freshly plated bacteria were grown overnight to logarithmic phase to an absorbance at 580 nm of 0.3 (Spectronic 88; Bausch and Lomb, Rochester, NY). Thigh infections with each of the pathogens were produced by injection of 0.1 mL of a 10^7.0^ CFU/mL inoculum into the thighs of isoflurane-anesthetized mice. Antibacterial therapy was initiated 2 hr post-infection. After 24 hr, animals were euthanized and the thighs were aseptically removed, homogenized, and plated for CFU determination. No-treatment controls were included in all experiments.

## Acknowledgments

The authors thank the University of Minnesota Genomics Center for metagenomic sequencing services and technical support. We thank Reed Stubbendieck (Oklahoma State University) for helpful discussions and assistance in biosynthetic gene cluster analyses. This work was supported by grants from the National Institutes of Health (U19AI142720 [C.R.C, L.R.K], NIAID F30AI169759 [S.Z], NCATS TL1TR002375 [S.Z], NIAID T32AI055397 [M.H.S], NIAMS F31AR079846 [M.H.S], U01AI125053 [N.S.]) and the Jarislowsky Foundation (C.R.C.). The funders had no role in study design, data collection and interpretation, or the decision to submit the work for publication. We declare that there are no conflicts of interest.

## Data availability

Raw metagenomic sequences generated for this work are deposited in the Short Read Archive under BioProject accession PRJNA1088473. All scripts and derived data necessary to replicate this work are available here: https://github.com/szelasko1/OralMDRO_paper and here for BGC analysis: https://github.com/reedstubbendieck/adt_bgcs.

