## Supplemental Figures and Tables for "Altered oral microbiota of drug-resistant organism carriers exhibit impaired gram-negative pathogen inhibition"

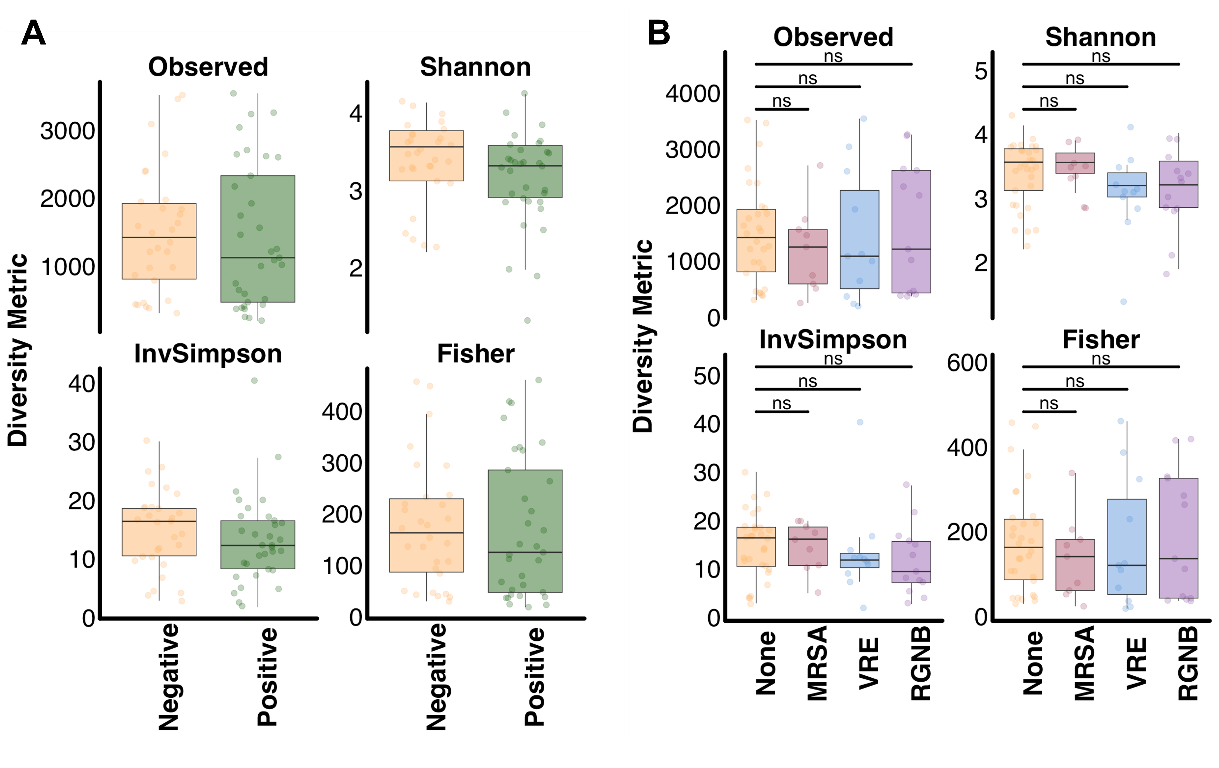

**Figure S1. Alpha Diversity of Oral Microbiomes from MDRO Carriers and Non-carriers.** Boxplot comparison of richness, Shannon, Inverse Simpson, and Fisher diversity metrics of oral microbiomes from (A) MDRO non-carriers (yellow) versus carriers (green) and (B) MDRO non-carriers (yellow) versus MRSA (red), VRE (blue), and RGNB (purple) carriers. Overlying data points representing diversity metric values have been jittered to avoid over-plotting. Significance value NS = p-value > 0.05, Mann–Whitney U-test.

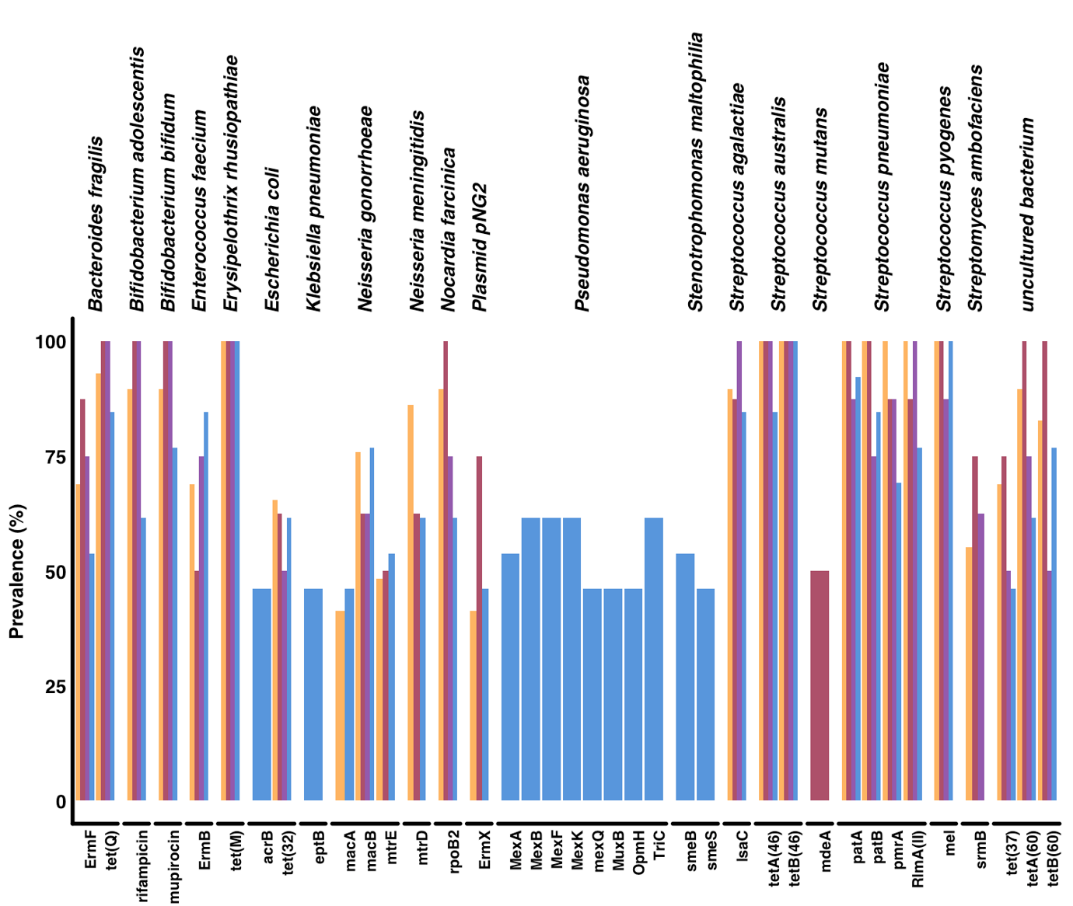

**Figure S2. Prevalence of AMR Genes between MDRO carriers and non-carriers**. Plot displays resistance genes with >40% prevalence across MDRO non-carriers (yellow), MRSA (red), VRE (purple), and RGNB (blue) carriers. Genes are grouped by pathogen of origin.

**Table S1. Beta Diversity of Oral Microbiomes from MDRO Carriers and Non-carriers.** ANOSIM R statistic and p-value with Benjamini-Hochberg correction comparisons of Bray-Curtis and Jaccard dissimilarity values between oral microbiomes of MDRO carriers and non-carriers (“None”). Permutations = 9999. Shotgun metagenomic sequencing reads were examined from microbiomes normalized to an equal sequencing depth. NS = not significant, ** = <0.01.

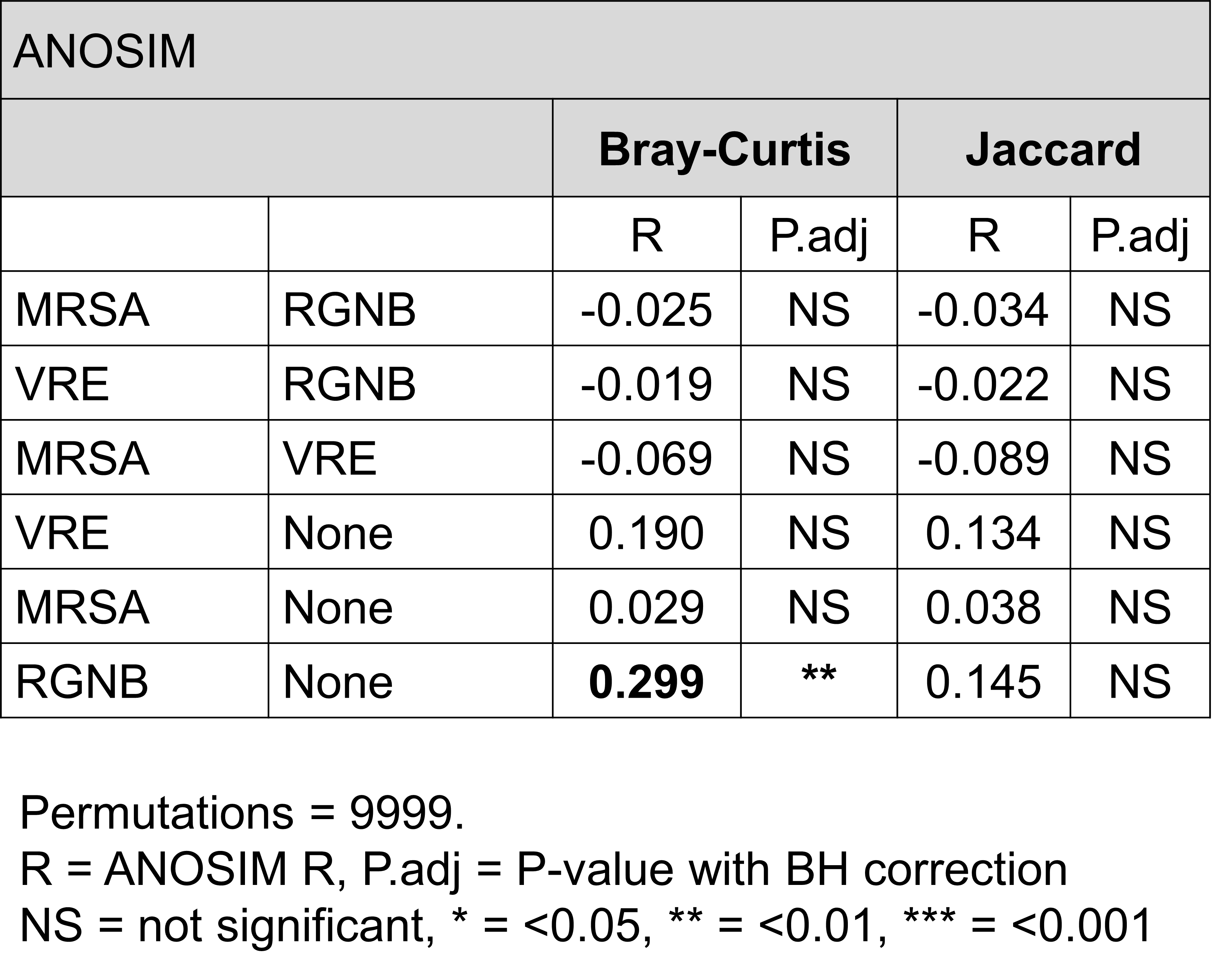

**Table S2. Differential Abundance Analysis Results.** List of species found to be differentially abundant between the oral microbiomes of RGNB carriers and non-carriers via edgeR analysis.

| **Species** | **logFC** | **logCPM** | **PValue** | **FDR** |
| --- | --- | --- | --- | --- |
| *Stenotrophomonas maltophilia* | -9.93 | 13.65 | 3.14E-29 | 1.32E-27 |
| *Serratia marcescens* | -11.99 | 13.40 | 3.56E-25 | 7.48E-24 |
| *Pseudomonas sp. SWI7* | -13.01 | 13.11 | 1.35E-23 | 1.89E-22 |
| *Pseudomonas fulva* | -13.14 | 14.16 | 1.87E-22 | 1.92E-21 |
| *Acinetobacter pittii* | -12.00 | 14.89 | 2.28E-22 | 1.92E-21 |
| *Acinetobacter johnsonii* | -10.58 | 15.05 | 1.30E-20 | 9.08E-20 |
| *Pseudomonas moraviensis* | -11.52 | 15.03 | 1.32E-17 | 7.93E-17 |
| *Pseudomonas atacamensis* | -11.63 | 15.62 | 9.66E-17 | 5.07E-16 |
| *Acinetobacter sp. NEB149* | -10.00 | 14.60 | 1.45E-14 | 6.77E-14 |
| *Klebsiella pneumoniae* | -8.24 | 13.63 | 3.52E-13 | 1.48E-12 |
| *Acinetobacter lwoffii* | -8.51 | 15.70 | 7.29E-12 | 2.78E-11 |
| *Limosilactobacillus fermentum* | -6.15 | 13.10 | 6.05E-08 | 2.12E-07 |
| *Streptococcus sp. HSISS2* | -3.94 | 14.61 | 9.98E-07 | 3.22E-06 |
| *Streptococcus sp. HSISS3* | -4.12 | 15.48 | 1.97E-06 | 5.91E-06 |
| *Streptococcus mitis* | 3.14 | 15.58 | 2.66E-05 | 7.46E-05 |
| *Streptococcus vestibularis* | -3.11 | 13.37 | 2.86E-05 | 7.50E-05 |
| *Streptococcus sp. SMGC_200* | 2.96 | 12.58 | 0.000154 | 0.000381 |
| *Streptococcus pneumoniae* | 2.38 | 14.04 | 0.00038 | 0.000887 |

**Table S3. Prevalence of AMR Genes between MDRO carriers and non-carriers.** Prevalence of genes across MDRO carriers and non-carriers.

| **ARG** | **Prev_all** | **Prev_none** | **Prev_MRSA** | **Prev_RGNB** | **Prev_VRE** | **Pathogen** |
| --- | --- | --- | --- | --- | --- | --- |
| tet(M) | 100.0 | 100.0 | 100.0 | 100.0 | 100.0 | *Erysipelothrix rhusiopathiae* |
| tetB(46) | 100.0 | 100.0 | 100.0 | 100.0 | 100.0 | *Streptococcus australis* |
| mel | 98.3 | 100.0 | 100.0 | 100.0 | 87.5 | *Streptococcus pyogenes* |
| patA | 96.6 | 100.0 | 100.0 | 92.3 | 87.5 | *Streptococcus pneumoniae* |
| tetA(46) | 96.6 | 100.0 | 100.0 | 84.6 | 100.0 | *Streptococcus australis* |
| patB | 93.1 | 100.0 | 100.0 | 84.6 | 75.0 | *Streptococcus pneumoniae* |
| RlmA(II) | 93.1 | 100.0 | 87.5 | 76.9 | 100.0 | *Streptococcus pneumoniae* |
| pmrA | 89.7 | 100.0 | 87.5 | 69.2 | 87.5 | *Streptococcus pneumoniae* |
| tet(Q) | 93.1 | 93.1 | 100.0 | 84.6 | 100.0 | *Bacteroides fragilis* |
| lsaC | 89.7 | 89.7 | 87.5 | 84.6 | 100.0 | *Streptococcus agalactiae* |
| Bifidobacterium bifidum ileS | 89.7 | 89.7 | 100.0 | 76.9 | 100.0 | *Bifidobacterium bifidum* |
| Bifidobacterium adolescentis rpoB | 86.2 | 89.7 | 100.0 | 61.5 | 100.0 | *Bifidobacterium adolescentis* |
| rpoB2 | 82.8 | 89.7 | 100.0 | 61.5 | 75.0 | *Nocardia farcinica* |
| tetA(60) | 82.8 | 89.7 | 100.0 | 61.5 | 75.0 | *uncultured bacterium* |
| mtrD | 70.7 | 86.2 | 62.5 | 61.5 | 37.5 | *Neisseria meningitidis* |
| tetB(60) | 79.3 | 82.8 | 100.0 | 76.9 | 50.0 | *uncultured bacterium* |
| macB | 72.4 | 75.9 | 62.5 | 76.9 | 62.5 | *Neisseria gonorrhoeae* |
| ErmB | 70.7 | 69.0 | 50.0 | 84.6 | 75.0 | *Enterococcus faecium* |
| ErmF | 69.0 | 69.0 | 87.5 | 53.8 | 75.0 | *Bacteroides fragilis* |
| tet(37) | 62.1 | 69.0 | 75.0 | 46.2 | 50.0 | *uncultured bacterium* |
| tet(32) | 62.1 | 65.5 | 62.5 | 61.5 | 50.0 | *Escherichia coli* |
| srmB | 55.2 | 55.2 | 75.0 | 38.5 | 62.5 | *Streptomyces ambofaciens* |
| mtrE | 46.6 | 48.3 | 50.0 | 53.8 | 25.0 | *Neisseria gonorrhoeae* |
| ErmX | 46.6 | 41.4 | 75.0 | 46.2 | 37.5 | *Plasmid pNG2* |
| macA | 41.4 | 41.4 | 37.5 | 46.2 | 37.5 | *Neisseria gonorrhoeae* |
| farB | 31.0 | 37.9 | 25.0 | 30.8 | 12.5 | *Neisseria meningitidis* |
| farA | 29.3 | 34.5 | 12.5 | 30.8 | 25.0 | *Neisseria meningitidis* |
| lmrD | 25.9 | 34.5 | 37.5 | 0.0 | 25.0 | *Lactococcus lactis* |
| mtrC | 31.0 | 31.0 | 25.0 | 38.5 | 25.0 | *Neisseria meningitidis* |
| tet(O/M/O) | 31.0 | 31.0 | 25.0 | 30.8 | 37.5 | *Campylobacter coli* |
| OprJ | 25.9 | 31.0 | 0.0 | 38.5 | 12.5 | *Pseudomonas aeruginosa* |
| hmrM | 20.7 | 31.0 | 12.5 | 7.7 | 12.5 | *Haemophilus influenzae* |
| mef(E) | 25.9 | 27.6 | 37.5 | 23.1 | 12.5 | *Streptococcus pneumoniae* |
| tlrC | 25.9 | 27.6 | 37.5 | 15.4 | 25.0 | *Streptomyces fradiae* |
| mdeA | 29.3 | 24.1 | 50.0 | 30.8 | 25.0 | *Streptococcus mutans* |
| oleB | 24.1 | 24.1 | 12.5 | 30.8 | 25.0 | *Streptomyces antibioticus* |
| msbA | 22.4 | 24.1 | 0.0 | 38.5 | 12.5 | *Escherichia coli* |
| adeF | 20.7 | 24.1 | 0.0 | 23.1 | 25.0 | *Acinetobacter baumannii* |
| tet(O) | 13.8 | 24.1 | 0.0 | 7.7 | 0.0 | *Campylobacter jejuni* |
| dfrE | 19.0 | 20.7 | 25.0 | 23.1 | 0.0 | *Enterococcus faecalis* |
| emrA | 19.0 | 20.7 | 0.0 | 23.1 | 25.0 | *Escherichia coli* |
| MexB | 27.6 | 17.2 | 12.5 | 61.5 | 25.0 | *Pseudomonas aeruginosa* |
| mexY | 13.8 | 17.2 | 0.0 | 15.4 | 12.5 | *Pseudomonas aeruginosa* |
| PC1 beta-lactamase (blaZ) | 13.8 | 17.2 | 12.5 | 7.7 | 12.5 | *Staphylococcus aureus* |
| arlR | 10.3 | 17.2 | 0.0 | 7.7 | 0.0 | *Staphylococcus aureus* |
| RanA | 10.3 | 17.2 | 12.5 | 0.0 | 0.0 | *Riemerella anatipestifer* |
| LpsA | 8.6 | 17.2 | 0.0 | 0.0 | 0.0 | *Haemophilus influenzae* |
| mexQ | 19.0 | 13.8 | 0.0 | 46.2 | 12.5 | *Pseudomonas aeruginosa* |
| tet(B) | 13.8 | 13.8 | 0.0 | 30.8 | 0.0 | *Gram-negative bacterium* |
| Erm(38) | 12.1 | 13.8 | 0.0 | 7.7 | 25.0 | *Mycolicibacterium smegmatis* |
| APH(6)-Id | 10.3 | 13.8 | 0.0 | 7.7 | 12.5 | *Pseudomonas aeruginosa* |
| RbpA | 10.3 | 13.8 | 0.0 | 7.7 | 12.5 | *Mycolicibacterium smegmatis* |
| dfrC | 10.3 | 13.8 | 12.5 | 0.0 | 12.5 | *Staphylococcus epidermidis* |
| ErmC | 8.6 | 13.8 | 12.5 | 0.0 | 0.0 | *Staphylococcus epidermidis* |
| TriC | 24.1 | 10.3 | 25.0 | 61.5 | 12.5 | *Pseudomonas aeruginosa* |
| MexK | 22.4 | 10.3 | 12.5 | 61.5 | 12.5 | *Pseudomonas aeruginosa* |
| MexF | 20.7 | 10.3 | 0.0 | 61.5 | 12.5 | *Pseudomonas aeruginosa* |
| MexI | 17.2 | 10.3 | 12.5 | 38.5 | 12.5 | *Pseudomonas aeruginosa* |
| msrA | 13.8 | 10.3 | 37.5 | 7.7 | 12.5 | *Staphylococcus epidermidis* |
| novA | 13.8 | 10.3 | 12.5 | 7.7 | 37.5 | *Streptomyces niveus* |
| acrD | 12.1 | 10.3 | 0.0 | 30.8 | 0.0 | *Escherichia coli* |
| OmpA | 12.1 | 10.3 | 0.0 | 23.1 | 12.5 | *Klebsiella pneumoniae* |
| AxyY | 12.1 | 10.3 | 25.0 | 15.4 | 0.0 | *Achromobacter insuavis* |
| MexD | 10.3 | 10.3 | 0.0 | 23.1 | 0.0 | *Pseudomonas aeruginosa* |
| MuxC | 10.3 | 10.3 | 0.0 | 15.4 | 12.5 | *Pseudomonas aeruginosa* |
| tet(35) | 10.3 | 10.3 | 12.5 | 7.7 | 12.5 | *Vibrio harveyi* |
| Corynebacterium striatum tetA | 8.6 | 10.3 | 0.0 | 7.7 | 12.5 | *Corynebacterium striatum* |
| tet(K) | 6.9 | 10.3 | 12.5 | 0.0 | 0.0 | *Staphylococcus aureus* |
| arnA | 13.8 | 6.9 | 0.0 | 38.5 | 12.5 | *Pseudomonas aeruginosa* |
| LptD | 13.8 | 6.9 | 12.5 | 30.8 | 12.5 | *Klebsiella pneumoniae* |
| smeE | 12.1 | 6.9 | 0.0 | 38.5 | 0.0 | *Stenotrophomonas maltophilia* |
| mdtC | 10.3 | 6.9 | 0.0 | 30.8 | 0.0 | *Escherichia coli* |
| mexN | 10.3 | 6.9 | 0.0 | 30.8 | 0.0 | *Pseudomonas aeruginosa* |
| ParR | 10.3 | 6.9 | 0.0 | 23.1 | 12.5 | *Pseudomonas aeruginosa* |
| TolC | 10.3 | 6.9 | 0.0 | 23.1 | 12.5 | *Escherichia coli* |
| catS | 8.6 | 6.9 | 0.0 | 15.4 | 12.5 | *Streptococcus pyogenes* |
| MexH | 8.6 | 6.9 | 12.5 | 15.4 | 0.0 | *Pseudomonas aeruginosa* |
| oleC | 8.6 | 6.9 | 12.5 | 15.4 | 0.0 | *Streptomyces antibioticus* |
| opmE | 8.6 | 6.9 | 0.0 | 15.4 | 12.5 | *Pseudomonas aeruginosa* |
| cprS | 6.9 | 6.9 | 0.0 | 15.4 | 0.0 | *Pseudomonas aeruginosa* |
| kdpD | 6.9 | 6.9 | 0.0 | 15.4 | 0.0 | *Staphylococcus aureus* |
| carA | 6.9 | 6.9 | 12.5 | 7.7 | 0.0 | *Streptomyces thermotolerans* |
| Staphylococcus aureus mupB | 6.9 | 6.9 | 0.0 | 7.7 | 12.5 | *Staphylococcus aureus* |
| lmrC | 6.9 | 6.9 | 25.0 | 0.0 | 0.0 | *Streptomyces lincolnensis* |
| norA | 5.2 | 6.9 | 0.0 | 7.7 | 0.0 | *Staphylococcus epidermidis* |
| norC | 5.2 | 6.9 | 0.0 | 7.7 | 0.0 | *Staphylococcus aureus* |
| sdrM | 5.2 | 6.9 | 0.0 | 7.7 | 0.0 | *Staphylococcus aureus* |
| sul2 | 5.2 | 6.9 | 0.0 | 7.7 | 0.0 | *Vibrio cholerae* |
| tet(A) | 5.2 | 6.9 | 0.0 | 7.7 | 0.0 | *Shigella sonnei* |
| tet(S) | 5.2 | 6.9 | 0.0 | 7.7 | 0.0 | *Listeria monocytogenes* |
| vanXY gene in vanG cluster | 5.2 | 6.9 | 0.0 | 7.7 | 0.0 | *Enterococcus faecalis* |
| cmx | 5.2 | 6.9 | 0.0 | 0.0 | 12.5 | *Corynebacterium striatum* |
| efrB | 5.2 | 6.9 | 0.0 | 0.0 | 12.5 | *Enterococcus faecium* |
| 23S rRNA (adenine(2058)-N(6))-methyltransferase Erm(A) | 3.4 | 6.9 | 0.0 | 0.0 | 0.0 | *Streptococcus pyogenes* |
| mdsC | 3.4 | 6.9 | 0.0 | 0.0 | 0.0 | *Salmonella enterica* |
| mecR1 | 3.4 | 6.9 | 0.0 | 0.0 | 0.0 | *Staphylococcus aureus* |
| tet(L) | 3.4 | 6.9 | 0.0 | 0.0 | 0.0 | *Geobacillus stearothermophilus* |
| tet(O/32/O) | 3.4 | 6.9 | 0.0 | 0.0 | 0.0 | *Clostridiaceae bacterium* |
| smeB | 19.0 | 3.4 | 25.0 | 53.8 | 12.5 | *Stenotrophomonas maltophilia* |
| MexA | 15.5 | 3.4 | 12.5 | 53.8 | 0.0 | *Pseudomonas aeruginosa* |
| OpmH | 15.5 | 3.4 | 0.0 | 46.2 | 25.0 | *Pseudomonas aeruginosa* |
| acrB | 13.8 | 3.4 | 12.5 | 46.2 | 0.0 | *Escherichia coli* |
| eptB | 13.8 | 3.4 | 0.0 | 46.2 | 12.5 | *Klebsiella pneumoniae* |
| MuxB | 13.8 | 3.4 | 0.0 | 46.2 | 12.5 | *Pseudomonas aeruginosa* |
| YajC | 13.8 | 3.4 | 12.5 | 38.5 | 12.5 | *Pseudomonas aeruginosa* |
| MexE | 12.1 | 3.4 | 25.0 | 30.8 | 0.0 | *Pseudomonas aeruginosa* |
| MexW | 12.1 | 3.4 | 12.5 | 30.8 | 12.5 | *Pseudomonas aeruginosa* |
| OprM | 10.3 | 3.4 | 0.0 | 38.5 | 0.0 | *Pseudomonas aeruginosa* |
| OprN | 10.3 | 3.4 | 0.0 | 38.5 | 0.0 | *Pseudomonas aeruginosa* |
| adeJ | 10.3 | 3.4 | 12.5 | 23.1 | 12.5 | *Acinetobacter baumannii* |
| OpmB | 8.6 | 3.4 | 0.0 | 30.8 | 0.0 | *Pseudomonas aeruginosa* |
| adeK | 8.6 | 3.4 | 12.5 | 23.1 | 0.0 | *Acinetobacter baumannii* |
| CRP | 8.6 | 3.4 | 0.0 | 23.1 | 12.5 | *Escherichia coli* |
| Escherichia coli mdfA | 8.6 | 3.4 | 12.5 | 15.4 | 12.5 | *Escherichia coli* |
| mgrA | 6.9 | 3.4 | 0.0 | 23.1 | 0.0 | *Staphylococcus aureus* |
| PmrF | 6.9 | 3.4 | 0.0 | 23.1 | 0.0 | *Escherichia coli* |
| basS | 6.9 | 3.4 | 12.5 | 15.4 | 0.0 | *Pseudomonas aeruginosa* |
| AcrF | 6.9 | 3.4 | 0.0 | 7.7 | 25.0 | *Escherichia coli* |
| APH(3')-Ia | 6.9 | 3.4 | 12.5 | 7.7 | 12.5 | *Serratia marcescens* |
| Staphylococcus aureus mupA | 6.9 | 3.4 | 25.0 | 0.0 | 12.5 | *Staphylococcus aureus* |
| APH(3')-IIb | 5.2 | 3.4 | 0.0 | 15.4 | 0.0 | *Pseudomonas aeruginosa* |
| efrA | 5.2 | 3.4 | 0.0 | 15.4 | 0.0 | *Enterococcus faecalis* |
| Enterobacter cloacae acrA | 5.2 | 3.4 | 0.0 | 15.4 | 0.0 | *Enterobacter cloacae* |
| arlS | 5.2 | 3.4 | 0.0 | 7.7 | 12.5 | *Staphylococcus aureus* |
| gadX | 5.2 | 3.4 | 12.5 | 7.7 | 0.0 | *Escherichia coli* |
| catA1 | 3.4 | 3.4 | 0.0 | 7.7 | 0.0 | *Escherichia coli* |
| lnuC | 3.4 | 3.4 | 0.0 | 7.7 | 0.0 | *Streptococcus agalactiae* |
| mdtP | 3.4 | 3.4 | 0.0 | 7.7 | 0.0 | *Escherichia coli* |
| mepA | 3.4 | 3.4 | 0.0 | 7.7 | 0.0 | *Staphylococcus aureus* |
| mepR | 3.4 | 3.4 | 0.0 | 7.7 | 0.0 | *Staphylococcus aureus* |
| pgpB | 3.4 | 3.4 | 0.0 | 7.7 | 0.0 | *Porphyromonas gingivalis* |
| poxtA | 3.4 | 3.4 | 0.0 | 7.7 | 0.0 | *Staphylococcus aureus* |
| Staphylococcus aureus LmrS | 3.4 | 3.4 | 0.0 | 7.7 | 0.0 | *Staphylococcus aureus* |
| Staphylococcus aureus norA | 3.4 | 3.4 | 0.0 | 7.7 | 0.0 | *Staphylococcus aureus* |
| tet(38) | 3.4 | 3.4 | 0.0 | 7.7 | 0.0 | *Staphylococcus aureus* |
| tet(C) | 3.4 | 3.4 | 0.0 | 7.7 | 0.0 | *Aeromonas salmonicida* |
| AAC(6')-Ie-APH(2'')-Ia bifunctional protein | 3.4 | 3.4 | 12.5 | 0.0 | 0.0 | *Staphylococcus aureus* |
| gadW | 3.4 | 3.4 | 12.5 | 0.0 | 0.0 | *Escherichia coli* |
| mdtO | 3.4 | 3.4 | 12.5 | 0.0 | 0.0 | *Escherichia coli* |
| OXA-85 | 3.4 | 3.4 | 12.5 | 0.0 | 0.0 | *Fusobacterium nucleatum* |
| TaeA | 3.4 | 3.4 | 0.0 | 0.0 | 12.5 | *Paenibacillus sp.* |
| aad(6) | 1.7 | 3.4 | 0.0 | 0.0 | 0.0 | *Streptococcus oralis* |
| APH(3'')-Ib | 1.7 | 3.4 | 0.0 | 0.0 | 0.0 | *Pseudomonas aeruginosa* |
| catQ | 1.7 | 3.4 | 0.0 | 0.0 | 0.0 | *Clostridium perfringens* |
| Clostridioides difficile cplR | 1.7 | 3.4 | 0.0 | 0.0 | 0.0 | *Clostridioides difficile* |
| emrK | 1.7 | 3.4 | 0.0 | 0.0 | 0.0 | *Escherichia coli* |
| emrY | 1.7 | 3.4 | 0.0 | 0.0 | 0.0 | *Escherichia coli* |
| EreD | 1.7 | 3.4 | 0.0 | 0.0 | 0.0 | *Riemerella anatipestifer* |
| Erm(36) | 1.7 | 3.4 | 0.0 | 0.0 | 0.0 | *Micrococcus luteus* |
| IreK | 1.7 | 3.4 | 0.0 | 0.0 | 0.0 | *Enterococcus faecalis* |
| lnuA | 1.7 | 3.4 | 0.0 | 0.0 | 0.0 | *Staphylococcus haemolyticus* |
| mdtE | 1.7 | 3.4 | 0.0 | 0.0 | 0.0 | *Escherichia coli* |
| mecA | 1.7 | 3.4 | 0.0 | 0.0 | 0.0 | *Staphylococcus aureus* |
| mphC | 1.7 | 3.4 | 0.0 | 0.0 | 0.0 | *Staphylococcus aureus* |
| mphE | 1.7 | 3.4 | 0.0 | 0.0 | 0.0 | *uncultured bacterium* |
| msrH | 1.7 | 3.4 | 0.0 | 0.0 | 0.0 | *Macrococcus canis* |
| PAM-1 | 1.7 | 3.4 | 0.0 | 0.0 | 0.0 | *Pseudomonas alcaligenes* |
| qacA | 1.7 | 3.4 | 0.0 | 0.0 | 0.0 | *Staphylococcus aureus* |
| SAT-4 | 1.7 | 3.4 | 0.0 | 0.0 | 0.0 | *Campylobacter coli* |
| tet(33) | 1.7 | 3.4 | 0.0 | 0.0 | 0.0 | *Corynebacterium glutamicum* |
| vanR gene in vanG cluster | 1.7 | 3.4 | 0.0 | 0.0 | 0.0 | *Enterococcus faecalis* |
| vanR gene in vanM cluster | 1.7 | 3.4 | 0.0 | 0.0 | 0.0 | *Enterococcus faecium* |
| smeF | 13.8 | 0.0 | 25.0 | 38.5 | 12.5 | *Stenotrophomonas maltophilia* |
| smeS | 10.3 | 0.0 | 0.0 | 46.2 | 0.0 | *Stenotrophomonas maltophilia* |
| Klebsiella pneumoniae KpnG | 10.3 | 0.0 | 0.0 | 38.5 | 12.5 | *Klebsiella sp.* |
| oqxB | 10.3 | 0.0 | 0.0 | 38.5 | 12.5 | *Escherichia coli* |
| Pseudomonas aeruginosa CpxR | 10.3 | 0.0 | 12.5 | 38.5 | 0.0 | *Pseudomonas aeruginosa* |
| bcr-1 | 10.3 | 0.0 | 12.5 | 30.8 | 12.5 | *Pseudomonas aeruginosa* |
| APH(9)-Ic | 8.6 | 0.0 | 0.0 | 38.5 | 0.0 | *Stenotrophomonas maltophilia* |
| ArnT | 8.6 | 0.0 | 0.0 | 30.8 | 12.5 | *Klebsiella pneumoniae* |
| bacA | 8.6 | 0.0 | 0.0 | 30.8 | 12.5 | *Escherichia coli* |
| ceoB | 8.6 | 0.0 | 0.0 | 30.8 | 12.5 | *Burkholderia cepacia* |
| mdtB | 8.6 | 0.0 | 0.0 | 30.8 | 12.5 | *Escherichia coli* |
| smeD | 8.6 | 0.0 | 0.0 | 30.8 | 12.5 | *Stenotrophomonas maltophilia* |
| TriA | 8.6 | 0.0 | 12.5 | 30.8 | 0.0 | *Pseudomonas aeruginosa* |
| cpxA | 8.6 | 0.0 | 0.0 | 23.1 | 25.0 | *Escherichia coli* |
| Klebsiella pneumoniae OmpK37 | 8.6 | 0.0 | 0.0 | 23.1 | 25.0 | *Klebsiella pneumoniae* |
| APH(3')-IIc | 6.9 | 0.0 | 0.0 | 30.8 | 0.0 | *Stenotrophomonas maltophilia* |
| cprR | 6.9 | 0.0 | 0.0 | 30.8 | 0.0 | *Pseudomonas aeruginosa* |
| MuxA | 6.9 | 0.0 | 0.0 | 30.8 | 0.0 | *Pseudomonas aeruginosa* |
| rsmA | 6.9 | 0.0 | 0.0 | 30.8 | 0.0 | *Pseudomonas aeruginosa* |
| smeA | 6.9 | 0.0 | 0.0 | 30.8 | 0.0 | *Stenotrophomonas maltophilia* |
| smeR | 6.9 | 0.0 | 0.0 | 30.8 | 0.0 | *Stenotrophomonas maltophilia* |
| TriB | 6.9 | 0.0 | 0.0 | 30.8 | 0.0 | *Pseudomonas aeruginosa* |
| H-NS | 6.9 | 0.0 | 0.0 | 23.1 | 12.5 | *Escherichia coli* |
| kdpE | 6.9 | 0.0 | 0.0 | 23.1 | 12.5 | *Escherichia coli* |
| MdtK | 6.9 | 0.0 | 0.0 | 23.1 | 12.5 | *Salmonella enterica* |
| mexP | 6.9 | 0.0 | 12.5 | 23.1 | 0.0 | *Pseudomonas aeruginosa* |
| oqxA | 6.9 | 0.0 | 0.0 | 23.1 | 12.5 | *Escherichia coli* |
| ParS | 6.9 | 0.0 | 12.5 | 23.1 | 0.0 | *Pseudomonas aeruginosa* |
| mdtF | 6.9 | 0.0 | 12.5 | 15.4 | 12.5 | *Escherichia coli* |
| emrR | 5.2 | 0.0 | 0.0 | 23.1 | 0.0 | *Escherichia coli* |
| Klebsiella pneumoniae acrA | 5.2 | 0.0 | 0.0 | 23.1 | 0.0 | *Klebsiella pneumoniae* |
| marA | 5.2 | 0.0 | 0.0 | 23.1 | 0.0 | *Escherichia coli* |
| mdtH | 5.2 | 0.0 | 0.0 | 23.1 | 0.0 | *Escherichia coli* |
| MexJ | 5.2 | 0.0 | 0.0 | 23.1 | 0.0 | *Pseudomonas aeruginosa* |
| MexL | 5.2 | 0.0 | 0.0 | 23.1 | 0.0 | *Pseudomonas aeruginosa* |
| PmpM | 5.2 | 0.0 | 0.0 | 23.1 | 0.0 | *Pseudomonas aeruginosa* |
| Pseudomonas aeruginosa catB7 | 5.2 | 0.0 | 0.0 | 23.1 | 0.0 | *Pseudomonas aeruginosa* |
| ramA | 5.2 | 0.0 | 0.0 | 23.1 | 0.0 | *Enterobacter cloacae* |
| smeC | 5.2 | 0.0 | 0.0 | 23.1 | 0.0 | *Stenotrophomonas maltophilia* |
| ugd | 5.2 | 0.0 | 0.0 | 23.1 | 0.0 | *Escherichia coli* |
| YojI | 5.2 | 0.0 | 0.0 | 23.1 | 0.0 | *Escherichia coli* |
| abeM | 5.2 | 0.0 | 0.0 | 15.4 | 12.5 | *Acinetobacter baumannii* |
| abeS | 5.2 | 0.0 | 12.5 | 15.4 | 0.0 | *Acinetobacter baumannii* |
| Acinetobacter baumannii AbaQ | 5.2 | 0.0 | 0.0 | 15.4 | 12.5 | *Acinetobacter baumannii* |
| ANT(3'')-IIa | 5.2 | 0.0 | 0.0 | 15.4 | 12.5 | *Acinetobacter baumannii* |
| eptA | 5.2 | 0.0 | 0.0 | 15.4 | 12.5 | *Escherichia coli* |
| mdtM | 5.2 | 0.0 | 12.5 | 15.4 | 0.0 | *Escherichia coli* |
| MdtQ | 5.2 | 0.0 | 0.0 | 15.4 | 12.5 | *Klebsiella pneumoniae* |
| MexC | 5.2 | 0.0 | 0.0 | 15.4 | 12.5 | *Pseudomonas aeruginosa* |
| OprA | 5.2 | 0.0 | 12.5 | 15.4 | 0.0 | *Pseudomonas aeruginosa* |
| rosA | 5.2 | 0.0 | 0.0 | 15.4 | 12.5 | *Yersinia enterocolitica* |
| AAC(6')-Ic | 3.4 | 0.0 | 0.0 | 15.4 | 0.0 | *Serratia marcescens* |
| AAC(6')-Iz | 3.4 | 0.0 | 0.0 | 15.4 | 0.0 | *Stenotrophomonas maltophilia* |
| Acinetobacter baumannii AmvA | 3.4 | 0.0 | 0.0 | 15.4 | 0.0 | *Acinetobacter baumannii* |
| adeB | 3.4 | 0.0 | 0.0 | 15.4 | 0.0 | *Acinetobacter baumannii* |
| adeG | 3.4 | 0.0 | 0.0 | 15.4 | 0.0 | *Acinetobacter baumannii* |
| adeH | 3.4 | 0.0 | 0.0 | 15.4 | 0.0 | *Acinetobacter baumannii* |
| adeI | 3.4 | 0.0 | 0.0 | 15.4 | 0.0 | *Acinetobacter baumannii* |
| adeL | 3.4 | 0.0 | 0.0 | 15.4 | 0.0 | *Acinetobacter baumannii* |
| adeN | 3.4 | 0.0 | 0.0 | 15.4 | 0.0 | *Acinetobacter baumannii* |
| adeS | 3.4 | 0.0 | 0.0 | 15.4 | 0.0 | *Acinetobacter baumannii* |
| FosA2 | 3.4 | 0.0 | 0.0 | 15.4 | 0.0 | *Enterobacter cloacae* |
| Klebsiella pneumoniae KpnE | 3.4 | 0.0 | 0.0 | 15.4 | 0.0 | *Klebsiella pneumoniae* |
| Klebsiella pneumoniae KpnF | 3.4 | 0.0 | 0.0 | 15.4 | 0.0 | *Klebsiella pneumoniae* |
| L1 beta-lactamase | 3.4 | 0.0 | 0.0 | 15.4 | 0.0 | *Stenotrophomonas maltophilia* |
| leuO | 3.4 | 0.0 | 0.0 | 15.4 | 0.0 | *Escherichia coli* |
| LpsB | 3.4 | 0.0 | 0.0 | 15.4 | 0.0 | *Acinetobacter baumannii* |
| mdtG | 3.4 | 0.0 | 0.0 | 15.4 | 0.0 | *Escherichia coli* |
| MexV | 3.4 | 0.0 | 0.0 | 15.4 | 0.0 | *Pseudomonas aeruginosa* |
| mexX | 3.4 | 0.0 | 0.0 | 15.4 | 0.0 | *Pseudomonas aeruginosa* |
| OpmD | 3.4 | 0.0 | 0.0 | 15.4 | 0.0 | *Pseudomonas aeruginosa* |
| Pseudomonas aeruginosa soxR | 3.4 | 0.0 | 0.0 | 15.4 | 0.0 | *Pseudomonas aeruginosa* |
| sdiA | 3.4 | 0.0 | 0.0 | 15.4 | 0.0 | *Salmonella enterica* |
| tet(41) | 3.4 | 0.0 | 0.0 | 15.4 | 0.0 | *Serratia marcescens* |
| rosB | 3.4 | 0.0 | 0.0 | 7.7 | 12.5 | *Yersinia enterocolitica* |
| aadA5 | 1.7 | 0.0 | 0.0 | 7.7 | 0.0 | *Escherichia coli* |
| AcrE | 1.7 | 0.0 | 0.0 | 7.7 | 0.0 | *Escherichia coli* |
| adeC | 1.7 | 0.0 | 0.0 | 7.7 | 0.0 | *Acinetobacter baumannii* |
| amrB | 1.7 | 0.0 | 0.0 | 7.7 | 0.0 | *Burkholderia pseudomallei* |
| ANT(4')-Ia | 1.7 | 0.0 | 0.0 | 7.7 | 0.0 | *Plasmid pTB913* |
| ANT(9)-Ia | 1.7 | 0.0 | 0.0 | 7.7 | 0.0 | *Staphylococcus aureus* |
| APH(3')-IIa | 1.7 | 0.0 | 0.0 | 7.7 | 0.0 | *Escherichia coli* |
| ArmR | 1.7 | 0.0 | 0.0 | 7.7 | 0.0 | *Pseudomonas aeruginosa* |
| AxyX | 1.7 | 0.0 | 0.0 | 7.7 | 0.0 | *Achromobacter insuavis* |
| ceoA | 1.7 | 0.0 | 0.0 | 7.7 | 0.0 | *Burkholderia cepacia* |
| CGA-1 | 1.7 | 0.0 | 0.0 | 7.7 | 0.0 | *Chryseobacterium gleum* |
| CGB-1 | 1.7 | 0.0 | 0.0 | 7.7 | 0.0 | *Chryseobacterium gleum* |
| CMY-157 | 1.7 | 0.0 | 0.0 | 7.7 | 0.0 | *Citrobacter sp.* |
| dfrA26 | 1.7 | 0.0 | 0.0 | 7.7 | 0.0 | *Escherichia coli* |
| ErmA | 1.7 | 0.0 | 0.0 | 7.7 | 0.0 | *Staphylococcus aureus* |
| FosA | 1.7 | 0.0 | 0.0 | 7.7 | 0.0 | *Pseudomonas aeruginosa* |
| GIL-1 | 1.7 | 0.0 | 0.0 | 7.7 | 0.0 | *Bacteria, Viruses,* |
| golS | 1.7 | 0.0 | 0.0 | 7.7 | 0.0 | *Salmonella enterica* |
| IND-9 | 1.7 | 0.0 | 0.0 | 7.7 | 0.0 | *Chryseobacterium indologenes* |
| LRA-10 | 1.7 | 0.0 | 0.0 | 7.7 | 0.0 | *uncultured bacterium* |
| LRA-13 | 1.7 | 0.0 | 0.0 | 7.7 | 0.0 | *uncultured bacterium* |
| MCR-7.1 | 1.7 | 0.0 | 0.0 | 7.7 | 0.0 | *Klebsiella pneumoniae* |
| mdsB | 1.7 | 0.0 | 0.0 | 7.7 | 0.0 | *Salmonella enterica* |
| mdtA | 1.7 | 0.0 | 0.0 | 7.7 | 0.0 | *Escherichia coli* |
| MexG | 1.7 | 0.0 | 0.0 | 7.7 | 0.0 | *Pseudomonas aeruginosa* |
| mexM | 1.7 | 0.0 | 0.0 | 7.7 | 0.0 | *Pseudomonas aeruginosa* |
| OprZ | 1.7 | 0.0 | 0.0 | 7.7 | 0.0 | *Achromobacter insuavis* |
| OXA-646 | 1.7 | 0.0 | 0.0 | 7.7 | 0.0 | *Acinetobacter lwoffii* |
| OXA-913 | 1.7 | 0.0 | 0.0 | 7.7 | 0.0 | *Pseudomonas* |
| Pseudomonas aeruginosa emrE | 1.7 | 0.0 | 0.0 | 7.7 | 0.0 | *Pseudomonas aeruginosa* |
| QnrB39 | 1.7 | 0.0 | 0.0 | 7.7 | 0.0 | *Citrobacter youngae* |
| sepA | 1.7 | 0.0 | 0.0 | 7.7 | 0.0 | *Staphylococcus* |
| Staphylococcus aureus FosB | 1.7 | 0.0 | 0.0 | 7.7 | 0.0 | *Staphylococcus aureus* |
| tet(W) | 1.7 | 0.0 | 0.0 | 7.7 | 0.0 | *Butyrivibrio fibrisolvens* |
| vanS gene in vanO cluster | 1.7 | 0.0 | 0.0 | 7.7 | 0.0 | *Rhodococcus hoagii* |
| Agrobacterium fabrum chloramphenicol acetyltransferase | 1.7 | 0.0 | 12.5 | 0.0 | 0.0 | *Agrobacterium fabrum* |
| catP | 1.7 | 0.0 | 12.5 | 0.0 | 0.0 | *Clostridium perfringens* |
| cmlA9 | 1.7 | 0.0 | 12.5 | 0.0 | 0.0 | *Salmonella enterica* |
| evgS | 1.7 | 0.0 | 12.5 | 0.0 | 0.0 | *Escherichia coli* |
| lsaA | 1.7 | 0.0 | 0.0 | 0.0 | 12.5 | *Enterococcus faecalis* |
| mdtN | 1.7 | 0.0 | 12.5 | 0.0 | 0.0 | *Escherichia coli* |
| Nocardia farcinica rox | 1.7 | 0.0 | 12.5 | 0.0 | 0.0 | *Nocardia farcinica* |
| tet(O/W/32/O) | 1.7 | 0.0 | 0.0 | 0.0 | 12.5 | *Lactobacillus johnsonii* |
| tet(T) | 1.7 | 0.0 | 0.0 | 0.0 | 12.5 | *Streptococcus pyogenes* |
| vanR gene in vanE cluster | 1.7 | 0.0 | 0.0 | 0.0 | 12.5 | *Enterococcus faecalis* |
| vanR gene in vanO cluster | 1.7 | 0.0 | 12.5 | 0.0 | 0.0 | *Rhodococcus hoagii* |

**Table S4. AMR Genes between RGNB carriers and MDRO non-carriers.** Genes identified to be differentially abundant between RGNB carriers and non-carriers used to create Figure 2C.

| **Gene** | **logFC** | **logCPM** | **PValue** | **FDR** | **log(FDR)** | **MDRO** |
| --- | --- | --- | --- | --- | --- | --- |
| RlmA(II) | -2.8089 | 16.7257 | 0.0072 | 0.0136 | 1.8652 | None |
| patA | -3.0313 | 17.5180 | 0.0031 | 0.0068 | 2.1675 | None |
| patB | -3.1339 | 17.6489 | 0.0023 | 0.0053 | 2.2798 | None |
| pmrA | -3.3856 | 16.6684 | 0.0018 | 0.0042 | 2.3774 | None |
| tet(K) | -3.8141 | 11.6035 | 0.0255 | 0.0444 | 1.3522 | None |
| hmrM | -4.1200 | 12.0562 | 0.0167 | 0.0297 | 1.5267 | None |
| cprS | 3.4936 | 11.6557 | 0.0420 | 0.0702 | 1.1538 | NA |
| basS | 3.2673 | 11.5480 | 0.0925 | 0.1408 | 0.8515 | NA |
| acrD | 3.0441 | 12.3459 | 0.0605 | 0.0968 | 1.0140 | NA |
| lnuC | 3.0388 | 11.2599 | 0.0422 | 0.0702 | 1.1538 | NA |
| MexH | 2.9853 | 11.5200 | 0.1049 | 0.1562 | 0.8062 | NA |
| APH(3')-IIb | 2.9017 | 11.3762 | 0.0646 | 0.1025 | 0.9891 | NA |
| rosB | 2.7438 | 11.1816 | 0.0559 | 0.0907 | 1.0423 | NA |
| sdiA | 2.6833 | 11.1040 | 0.0559 | 0.0907 | 1.0423 | NA |
| PmrF | 2.6089 | 11.3979 | 0.1978 | 0.2670 | 0.5735 | NA |
| mepR | 2.4515 | 11.1429 | 0.0595 | 0.0960 | 1.0179 | NA |
| ParR | 2.4292 | 11.4401 | 0.2068 | 0.2775 | 0.5568 | NA |
| catA1 | 2.4229 | 11.1195 | 0.0307 | 0.0522 | 1.2823 | NA |
| pgpB | 2.2573 | 11.8904 | 0.1349 | 0.1943 | 0.7116 | NA |
| mepA | 2.1339 | 11.1903 | 0.0833 | 0.1286 | 0.8909 | NA |
| tet(38) | 2.1108 | 11.2570 | 0.0911 | 0.1396 | 0.8551 | NA |
| poxtA | 2.0998 | 11.1695 | 0.1819 | 0.2502 | 0.6017 | NA |
| norC | 2.0991 | 11.2870 | 0.1775 | 0.2490 | 0.6038 | NA |
| P. aeruginosa soxR | 2.0124 | 11.0603 | 0.1101 | 0.1619 | 0.7909 | NA |
| srmB | 1.9698 | 12.0796 | 0.0995 | 0.1502 | 0.8232 | NA |
| catS | 1.9370 | 11.3477 | 0.1045 | 0.1562 | 0.8062 | NA |
| Staphylococcus aureus norA | 1.7998 | 11.1377 | 0.1817 | 0.2502 | 0.6017 | NA |
| efrA | 1.7749 | 11.0757 | 0.1101 | 0.1619 | 0.7909 | NA |
| Staphylococcus aureus LmrS | 1.7721 | 11.1769 | 0.1976 | 0.2670 | 0.5735 | NA |
| MuxC | 1.7608 | 12.4509 | 0.2499 | 0.3332 | 0.4773 | NA |
| tet(S) | 1.6915 | 11.1951 | 0.1977 | 0.2670 | 0.5735 | NA |
| ErmF | 1.6409 | 14.3683 | 0.0821 | 0.1276 | 0.8941 | NA |
| oleC | 1.5533 | 11.1839 | 0.2809 | 0.3625 | 0.4407 | NA |
| farA | 1.5078 | 11.8848 | 0.1728 | 0.2456 | 0.6097 | NA |
| msrA | 1.4953 | 11.7005 | 0.3893 | 0.4860 | 0.3134 | NA |
| tet(M) | 1.3885 | 18.1330 | 0.0669 | 0.1055 | 0.9769 | NA |
| Erm(38) | 1.3407 | 11.4769 | 0.3598 | 0.4571 | 0.3400 | NA |
| tlrC | 1.3137 | 11.6662 | 0.3641 | 0.4599 | 0.3374 | NA |
| Bifidobacterium bifidum ileS | 1.2546 | 14.4514 | 0.1761 | 0.2486 | 0.6044 | NA |
| rpoB2 | 1.2193 | 14.8891 | 0.1802 | 0.2502 | 0.6017 | NA |
| mtrE | 1.2090 | 12.4288 | 0.2819 | 0.3625 | 0.4407 | NA |
| tetA(46) | 1.2023 | 16.2676 | 0.1326 | 0.1923 | 0.7161 | NA |
| tetB(46) | 1.1628 | 16.2853 | 0.1447 | 0.2069 | 0.6841 | NA |
| tet(C) | 1.1625 | 11.1585 | 0.6153 | 0.7223 | 0.1413 | NA |
| Staphylococcus aureus mupB | 1.0543 | 11.1211 | 0.2731 | 0.3554 | 0.4493 | NA |
| oleB | 1.0532 | 11.5919 | 0.4627 | 0.5647 | 0.2482 | NA |
| sdrM | 0.9765 | 11.0950 | 0.7384 | 0.8350 | 0.0783 | NA |
| gadX | 0.8895 | 11.1337 | 0.4481 | 0.5499 | 0.2597 | NA |
| Bifidobacterium adolescentis rpoB | 0.8840 | 16.0089 | 0.2670 | 0.3495 | 0.4565 | NA |
| arlS | 0.8760 | 11.0623 | 0.9961 | 1.0000 | 0.0000 | NA |
| tet(Q) | 0.8466 | 17.3342 | 0.2668 | 0.3495 | 0.4565 | NA |
| Corynebacterium striatum tetA | 0.8306 | 11.4286 | 0.7299 | 0.8298 | 0.0810 | NA |
| tet(32) | 0.7409 | 13.3396 | 0.4740 | 0.5753 | 0.2401 | NA |
| tet(B) | 0.7400 | 11.8361 | 0.4781 | 0.5769 | 0.2389 | NA |
| sul2 | 0.7046 | 11.1361 | 0.6153 | 0.7223 | 0.1413 | NA |
| APH(3')-Ia | 0.6627 | 11.2686 | 1.0000 | 1.0000 | 0.0000 | NA |
| mel | 0.6175 | 17.8775 | 0.4068 | 0.5021 | 0.2992 | NA |
| tet(A) | 0.5986 | 11.0748 | 1.0000 | 1.0000 | 0.0000 | NA |
| carA | 0.5788 | 11.2572 | 1.0000 | 1.0000 | 0.0000 | NA |
| farB | 0.5344 | 11.5026 | 0.7981 | 0.8886 | 0.0513 | NA |
| mtrD | 0.5097 | 14.4782 | 0.5617 | 0.6741 | 0.1713 | NA |
| macA | 0.4869 | 11.8546 | 0.6938 | 0.7929 | 0.1008 | NA |
| dfrE | 0.4717 | 11.3280 | 0.6606 | 0.7672 | 0.1151 | NA |
| mtrC | 0.4530 | 12.1665 | 0.6856 | 0.7915 | 0.1015 | NA |
| macB | 0.3476 | 14.1131 | 0.6889 | 0.7915 | 0.1015 | NA |
| mdtP | 0.1703 | 11.2100 | 1.0000 | 1.0000 | 0.0000 | NA |
| norA | 0.1487 | 11.2629 | 1.0000 | 1.0000 | 0.0000 | NA |
| novA | 0.1110 | 11.2442 | 1.0000 | 1.0000 | 0.0000 | NA |
| tet(O/M/O) | 0.0534 | 12.9182 | 0.8802 | 0.9700 | 0.0132 | NA |
| tet(35) | -0.1180 | 11.1150 | 1.0000 | 1.0000 | 0.0000 | NA |
| OXA-85 | -0.2677 | 11.0257 | 1.0000 | 1.0000 | 0.0000 | NA |
| TaeA | -0.3955 | 11.0411 | 1.0000 | 1.0000 | 0.0000 | NA |
| RbpA | -0.4197 | 11.1486 | 1.0000 | 1.0000 | 0.0000 | NA |
| mef(E) | -0.4397 | 11.8057 | 0.8517 | 0.9434 | 0.0253 | NA |
| mecR1 | -0.4960 | 11.0235 | 1.0000 | 1.0000 | 0.0000 | NA |
| tet(O) | -0.5721 | 11.3319 | 0.6257 | 0.7305 | 0.1364 | NA |
| AAC(6')-Ie-APH(2'')-Ia bifunctional protein | -0.6063 | 11.1092 | 1.0000 | 1.0000 | 0.0000 | NA |
| Staphylococcus aureus mupA | -0.6089 | 11.1428 | 1.0000 | 1.0000 | 0.0000 | NA |
| cmx | -0.6214 | 11.0994 | 1.0000 | 1.0000 | 0.0000 | NA |
| APH(6)-Id | -0.6231 | 11.5617 | 0.8902 | 0.9761 | 0.0105 | NA |
| mdsC | -0.7121 | 11.0433 | 1.0000 | 1.0000 | 0.0000 | NA |
| lmrC | -0.7941 | 11.1286 | 1.0000 | 1.0000 | 0.0000 | NA |
| efrB | -0.7995 | 11.0519 | 1.0000 | 1.0000 | 0.0000 | NA |
| tet(L) | -0.9285 | 11.0506 | 1.0000 | 1.0000 | 0.0000 | NA |
| arlR | -0.9517 | 11.2294 | 0.4043 | 0.5018 | 0.2994 | NA |
| mdtO | -1.1609 | 11.1675 | 0.5911 | 0.7020 | 0.1537 | NA |
| RanA | -1.1732 | 11.0908 | 0.7587 | 0.8491 | 0.0710 | NA |
| 23S rRNA (adenine(2058)-N(6))-methyltransferase Erm(A) | -1.2548 | 11.0788 | 0.7587 | 0.8491 | 0.0710 | NA |
| gadW | -1.2770 | 11.2076 | 0.3843 | 0.4826 | 0.3164 | NA |
| PC1 beta-lactamase (blaZ) | -1.3295 | 11.4956 | 0.3226 | 0.4124 | 0.3847 | NA |
| dfrC | -1.4332 | 11.1835 | 0.5915 | 0.7020 | 0.1537 | NA |
| ErmX | -1.6507 | 13.9522 | 0.2526 | 0.3347 | 0.4753 | NA |
| tet(O/32/O) | -2.5107 | 11.2784 | 0.1316 | 0.1920 | 0.7167 | NA |
| ErmC | -3.0219 | 11.4287 | 0.0769 | 0.1204 | 0.9193 | NA |
| LpsA | -3.4022 | 11.4084 | 0.0496 | 0.0818 | 1.0872 | NA |
| lmrD | -3.6717 | 11.6816 | 0.0309 | 0.0522 | 1.2823 | NA |
| adeJ | 9.1574 | 14.6834 | 1.72E-13 | 3.71E-11 | 10.4310 | RGNB |
| oqxB | 8.6637 | 13.7844 | 6.83E-12 | 5.13E-10 | 9.2899 | RGNB |
| acrB | 8.5640 | 13.9357 | 7.13E-12 | 5.13E-10 | 9.2899 | RGNB |
| Klebsiella pneumoniae OmpK37 | 7.7099 | 13.1217 | 6.56E-09 | 3.54E-07 | 6.4506 | RGNB |
| abeM | 7.6008 | 12.9168 | 2.85E-08 | 1.23E-06 | 5.9097 | RGNB |
| Acinetobacter baumannii AmvA | 7.5049 | 12.8173 | 5.26E-08 | 1.89E-06 | 5.7227 | RGNB |
| adeK | 7.4565 | 13.2735 | 2.09E-07 | 3.22E-06 | 5.4918 | RGNB |
| ANT(3'')-IIa | 7.4041 | 12.7658 | 6.72E-08 | 2.04E-06 | 5.6902 | RGNB |
| Acinetobacter baumannii AbaQ | 7.3245 | 12.7543 | 8.50E-08 | 2.04E-06 | 5.6902 | RGNB |
| adeL | 7.2391 | 12.6266 | 9.60E-08 | 2.07E-06 | 5.6833 | RGNB |
| LpsB | 7.0786 | 12.5142 | 1.39E-07 | 2.72E-06 | 5.5652 | RGNB |
| Pseudomonas aeruginosa CpxR | 7.0165 | 12.4880 | 1.53E-07 | 2.75E-06 | 5.5613 | RGNB |
| adeH | 6.9168 | 12.4105 | 2.01E-07 | 3.22E-06 | 5.4918 | RGNB |
| LptD | 6.8339 | 12.9743 | 7.36E-07 | 6.48E-06 | 5.1881 | RGNB |
| adeI | 6.8248 | 12.3521 | 2.35E-07 | 3.39E-06 | 5.4704 | RGNB |
| mdtB | 6.8095 | 12.4274 | 2.85E-07 | 3.85E-06 | 5.4145 | RGNB |
| MexW | 6.7847 | 13.3115 | 2.49E-05 | 0.0001 | 3.9942 | RGNB |
| MexK | 6.6594 | 13.7524 | 1.63E-06 | 1.13E-05 | 4.9451 | RGNB |
| adeG | 6.6277 | 12.2289 | 3.39E-07 | 4.31E-06 | 5.3660 | RGNB |
| Enterobacter cloacae acrA | 6.6217 | 12.6394 | 9.51E-05 | 0.0003 | 3.5326 | RGNB |
| smeF | 6.5777 | 12.2213 | 3.64E-07 | 4.37E-06 | 5.3594 | RGNB |
| MuxB | 6.5761 | 13.1943 | 6.02E-05 | 0.0002 | 3.6864 | RGNB |
| ArnT | 6.5372 | 12.3503 | 5.62E-07 | 5.91E-06 | 5.2288 | RGNB |
| CRP | 6.5277 | 12.4645 | 1.12E-06 | 8.68E-06 | 5.0617 | RGNB |
| adeF | 6.5257 | 13.6850 | 7.94E-08 | 2.04E-06 | 5.6902 | RGNB |
| Klebsiella pneumoniae KpnG | 6.5047 | 12.3174 | 5.74E-07 | 5.91E-06 | 5.2288 | RGNB |
| smeD | 6.4089 | 12.1348 | 5.34E-07 | 5.91E-06 | 5.2288 | RGNB |
| cpxA | 6.3159 | 12.1326 | 7.39E-07 | 6.48E-06 | 5.1881 | RGNB |
| OmpA | 6.2577 | 12.8493 | 8.48E-05 | 0.0003 | 3.5735 | RGNB |
| H-NS | 6.2460 | 12.0470 | 7.61E-07 | 6.48E-06 | 5.1881 | RGNB |
| emrR | 6.2370 | 11.9755 | 7.21E-07 | 6.48E-06 | 5.1881 | RGNB |
| oqxA | 6.2009 | 11.9816 | 7.80E-07 | 6.48E-06 | 5.1881 | RGNB |
| smeB | 6.1288 | 13.3190 | 0.0001 | 0.0003 | 3.4592 | RGNB |
| MexA | 6.1044 | 12.3241 | 4.02E-05 | 0.0001 | 3.8540 | RGNB |
| smeS | 6.0720 | 11.9202 | 1.04E-06 | 8.34E-06 | 5.0787 | RGNB |
| mdtC | 6.0356 | 13.0467 | 0.0002 | 0.0005 | 3.3063 | RGNB |
| YojI | 6.0194 | 11.8807 | 1.37E-06 | 9.84E-06 | 5.0069 | RGNB |
| smeC | 6.0158 | 11.9096 | 1.28E-06 | 9.56E-06 | 5.0194 | RGNB |
| MexF | 5.9853 | 14.1079 | 7.71E-06 | 4.27E-05 | 4.3698 | RGNB |
| Klebsiella pneumoniae acrA | 5.9387 | 11.8533 | 1.70E-06 | 1.14E-05 | 4.9415 | RGNB |
| PmpM | 5.9377 | 11.8418 | 1.82E-06 | 1.19E-05 | 4.9231 | RGNB |
| eptB | 5.9177 | 12.9063 | 0.0006 | 0.0014 | 2.8427 | RGNB |
| MdtQ | 5.8622 | 11.9362 | 2.39E-06 | 1.52E-05 | 4.8184 | RGNB |
| arnA | 5.8453 | 12.4493 | 0.0002 | 0.0006 | 3.1947 | RGNB |
| smeR | 5.7887 | 11.7675 | 3.16E-06 | 1.90E-05 | 4.7207 | RGNB |
| TriA | 5.7830 | 11.7515 | 3.44E-06 | 2.01E-05 | 4.6976 | RGNB |
| APH(9)-Ic | 5.7302 | 11.7722 | 3.17E-06 | 1.90E-05 | 4.7207 | RGNB |
| smeE | 5.6837 | 13.5200 | 0.0001 | 0.0004 | 3.4278 | RGNB |
| MuxA | 5.6445 | 11.7015 | 4.89E-06 | 2.78E-05 | 4.5564 | RGNB |
| AAC(6')-Ic | 5.5129 | 11.6738 | 8.03E-06 | 4.34E-05 | 4.3630 | RGNB |
| marA | 5.4980 | 11.6401 | 8.83E-06 | 4.45E-05 | 4.3512 | RGNB |
| ramA | 5.4593 | 11.6153 | 1.23E-05 | 5.79E-05 | 4.2377 | RGNB |
| smeA | 5.4372 | 11.6523 | 8.87E-06 | 4.45E-05 | 4.3512 | RGNB |
| mdtH | 5.4342 | 11.6435 | 8.84E-06 | 4.45E-05 | 4.3512 | RGNB |
| adeB | 5.3920 | 11.6314 | 1.10E-05 | 5.40E-05 | 4.2677 | RGNB |
| tet(41) | 5.3884 | 11.6263 | 1.23E-05 | 5.79E-05 | 4.2377 | RGNB |
| eptA | 5.2592 | 11.6440 | 1.78E-05 | 8.20E-05 | 4.0862 | RGNB |
| bacA | 5.2485 | 11.7452 | 1.85E-05 | 8.31E-05 | 4.0804 | RGNB |
| Escherichia coli mdfA | 5.2185 | 11.9030 | 0.0001 | 0.0004 | 3.4278 | RGNB |
| mdtM | 5.2153 | 11.6004 | 1.99E-05 | 8.61E-05 | 4.0649 | RGNB |
| MexJ | 5.2143 | 11.5187 | 2.90E-05 | 0.0001 | 3.9362 | RGNB |
| mdeA | 5.2043 | 13.6047 | 3.97E-05 | 0.0001 | 3.8540 | RGNB |
| adeS | 5.1986 | 11.5623 | 1.96E-05 | 8.61E-05 | 4.0649 | RGNB |
| abeS | 5.1928 | 11.5902 | 2.26E-05 | 9.59E-05 | 4.0184 | RGNB |
| OpmD | 5.1864 | 11.5145 | 3.34E-05 | 0.0001 | 3.8979 | RGNB |
| YajC | 5.1837 | 11.7928 | 2.47E-05 | 0.0001 | 3.9942 | RGNB |
| ceoB | 5.1724 | 11.5700 | 2.95E-05 | 0.0001 | 3.9362 | RGNB |
| Pseudomonas aeruginosa catB7 | 5.1503 | 11.5017 | 3.86E-05 | 0.0001 | 3.8575 | RGNB |
| OprM | 5.0870 | 12.2053 | 0.0031 | 0.0067 | 2.1709 | RGNB |
| mdtF | 5.0855 | 11.5875 | 3.42E-05 | 0.0001 | 3.8948 | RGNB |
| MexB | 5.0584 | 14.0328 | 7.24E-05 | 0.0002 | 3.6318 | RGNB |
| TriB | 5.0364 | 11.4855 | 3.85E-05 | 0.0001 | 3.8575 | RGNB |
| ParS | 5.0296 | 11.5126 | 3.34E-05 | 0.0001 | 3.8979 | RGNB |
| mexN | 4.9323 | 12.1944 | 0.0037 | 0.0079 | 2.1051 | RGNB |
| MexE | 4.9122 | 12.0772 | 0.0011 | 0.0025 | 2.5976 | RGNB |
| APH(3')-IIc | 4.8836 | 11.4513 | 6.10E-05 | 0.0002 | 3.6864 | RGNB |
| mexP | 4.8644 | 11.4483 | 7.20E-05 | 0.0002 | 3.6318 | RGNB |
| OprN | 4.8456 | 11.6708 | 0.0003 | 0.0009 | 3.0501 | RGNB |
| mexX | 4.8153 | 11.4360 | 7.18E-05 | 0.0002 | 3.6318 | RGNB |
| adeN | 4.8029 | 11.4381 | 8.53E-05 | 0.0003 | 3.5735 | RGNB |
| AcrF | 4.6928 | 12.1472 | 0.0072 | 0.0136 | 1.8652 | RGNB |
| Klebsiella pneumoniae KpnF | 4.6454 | 11.3814 | 0.0002 | 0.0005 | 3.2989 | RGNB |
| MdtK | 4.6160 | 11.4489 | 0.0001 | 0.0004 | 3.3693 | RGNB |
| cprR | 4.6138 | 11.3834 | 0.0001 | 0.0004 | 3.3693 | RGNB |
| TolC | 4.6011 | 12.2301 | 0.0059 | 0.0119 | 1.9228 | RGNB |
| mexQ | 4.5951 | 12.5395 | 0.0028 | 0.0062 | 2.2093 | RGNB |
| MexV | 4.5544 | 11.3612 | 0.0002 | 0.0006 | 3.2160 | RGNB |
| bcr-1 | 4.5023 | 11.3784 | 0.0003 | 0.0009 | 3.0508 | RGNB |
| TriC | 4.4763 | 13.7051 | 0.0007 | 0.0018 | 2.7543 | RGNB |
| MexC | 4.4445 | 11.3536 | 0.0003 | 0.0009 | 3.0508 | RGNB |
| OpmH | 4.2956 | 12.6451 | 0.0110 | 0.0201 | 1.6965 | RGNB |
| MexL | 4.2772 | 11.3047 | 0.0005 | 0.0014 | 2.8600 | RGNB |
| mexY | 4.2487 | 12.2136 | 0.0015 | 0.0036 | 2.4455 | RGNB |
| OprA | 4.2140 | 11.3221 | 0.0005 | 0.0014 | 2.8600 | RGNB |
| Klebsiella pneumoniae KpnE | 4.1484 | 11.2864 | 0.0007 | 0.0017 | 2.7591 | RGNB |
| OpmB | 4.1018 | 11.8764 | 0.0060 | 0.0120 | 1.9222 | RGNB |
| rosA | 3.9613 | 11.3557 | 0.0013 | 0.0029 | 2.5309 | RGNB |
| rsmA | 3.9096 | 11.2433 | 0.0012 | 0.0029 | 2.5312 | RGNB |
| ugd | 3.8287 | 11.2315 | 0.0023 | 0.0053 | 2.2798 | RGNB |
| leuO | 3.8213 | 11.2312 | 0.0048 | 0.0099 | 2.0036 | RGNB |
| MexD | 3.6375 | 12.2754 | 0.0094 | 0.0176 | 1.7549 | RGNB |
| AAC(6')-Iz | 3.6258 | 11.2008 | 0.0033 | 0.0071 | 2.1513 | RGNB |
| mdtG | 3.5878 | 11.1955 | 0.0048 | 0.0099 | 2.0036 | RGNB |
| kdpE | 3.5786 | 11.2545 | 0.0048 | 0.0099 | 2.0036 | RGNB |
| msbA | 3.5617 | 12.3700 | 0.0138 | 0.0248 | 1.6048 | RGNB |
| tetB(60) | 3.5046 | 14.3733 | 0.0003 | 0.0008 | 3.1160 | RGNB |
| AxyY | 3.3579 | 11.3498 | 0.0066 | 0.0131 | 1.8841 | RGNB |
| MexI | 3.3492 | 12.9372 | 0.0202 | 0.0356 | 1.4491 | RGNB |
| FosA2 | 3.3149 | 11.1729 | 0.0111 | 0.0202 | 1.6954 | RGNB |
| L1 beta-lactamase | 3.3005 | 11.1578 | 0.0072 | 0.0136 | 1.8652 | RGNB |
| kdpD | 3.2918 | 11.5543 | 0.0029 | 0.0065 | 2.1871 | RGNB |
| tetA(60) | 3.2841 | 14.9270 | 0.0003 | 0.0008 | 3.0809 | RGNB |
| mgrA | 3.2294 | 11.1984 | 0.0072 | 0.0136 | 1.8652 | RGNB |
| opmE | 3.2070 | 11.4905 | 0.0291 | 0.0498 | 1.3024 | RGNB |
| emrA | 3.1509 | 11.8780 | 0.0074 | 0.0139 | 1.8568 | RGNB |
| OprJ | 3.0286 | 11.8272 | 0.0068 | 0.0133 | 1.8769 | RGNB |
| vanXY gene in vanG cluster | 2.9310 | 11.2483 | 0.0184 | 0.0325 | 1.4878 | RGNB |
| ErmB | 2.6785 | 14.6290 | 0.0050 | 0.0102 | 1.9922 | RGNB |
| lsaC | 2.2181 | 15.4286 | 0.0099 | 0.0182 | 1.7389 | RGNB |
| tet(37) | 2.1987 | 13.9642 | 0.0263 | 0.0455 | 1.3422 | RGNB |
